# Spatiotemporal structure of sensory-evoked and spontaneous activity revealed by mesoscale imaging in anesthetized and awake mice

**DOI:** 10.1101/2020.05.22.111021

**Authors:** Navvab Afrashteh, Samsoon Inayat, Edgar Bermudez Contreras, Artur Luczak, Bruce L. McNaughton, Majid H. Mohajerani

**Author notes:** Corresponding author: Majid H. Mohajerani. These authors contributed equally (Navvab Afrashteh, Samsoon Inayat, Edgar Bermudez Contreras).

## Abstract

Brain activity propagates across the cortex in diverse spatiotemporal patterns, both as a response to sensory stimulation and during spontaneous activity. Despite been extensively studied, the relationship between the characteristics of such patterns during spontaneous and evoked activity is not completely understood. To investigate this relationship, we compared visual, auditory, and tactile evoked activity patterns elicited with different stimulus strengths and spontaneous activity motifs in lightly anesthetized and awake mice using mesoscale wide-field voltage-sensitive dye and glutamate imaging respectively. The characteristics of cortical activity that we compared include amplitude, speed, direction, and complexity of propagation trajectories in spontaneous and evoked activity patterns. We found that the complexity of the propagation trajectories of spontaneous activity, quantified as their fractal dimension, is higher than the one from sensory evoked responses. Moreover, the speed and direction of propagation, are modulated by the amplitude during both, spontaneous and evoked activity. Finally, we found that spontaneous activity had similar amplitude and speed when compared to evoked activity elicited with low stimulus strengths. However, this similarity gradually decreased when the strength of stimuli eliciting evoked responses increased. Altogether, these findings are consistent with the fact that even primary sensory areas receive widespread inputs from other cortical regions, and that, during rest, the cortex tends to reactivate traces of complex, multi-sensory experiences that may have occurred in a range of different behavioural contexts.

## 1 Introduction

Much of the development of our current understanding of sensory processing comes from the study of evoked activity and the stimulus-response relationships at different stages in the nervous system (Seung and Sompolinsky, 1993; Butts and Goldman, 2006; Jones and Smith, 2014). However, in the absence of stimuli the cortex remains active, even in the primary sensory areas. Until not long ago, spontaneous activity used to be regarded by some as ‘noise’ (Parga and Abbott, 2007; Faisal et al., 2008; Stringer et al., 2016). However, this view has been steadily replaced by the idea that spontaneous activity is crucial for the understanding of cortical function (McCormick, 1999; Ringach, 2009; Raichle, 2010; Deco et al., 2013).

Despite significant advances in the understanding of cortical processing underlying both spontaneous and evoked activity, their relationship and interactions remain under debate. In some studies, the spatiotemporal patterns of evoked and spontaneous activity are reported to be similar (Hoffman and McNaughton, 2002; Kenet et al., 2003; Han et al., 2008; Luczak et al., 2009; Berkes et al., 2011), while in others they are reported to have remarkable differences (Stringer et al., 2019). For example, at the microcircuit level, Luczak and colleagues proposed that the temporal patterns of tone-evoked spiking activity occur as part of a larger set of patterns produced during spontaneous activity recorded using silicon probes within auditory cortex in anesthetized rats (Luczak et al., 2009; Luczak et al., 2015). In contrast, more recent studies using large-scale high-density optical imaging as well as silicon probes to record the single-unit activity of large neuronal populations, report that the patterns of activity from evoked responses belong to a different space from the ones in spontaneous activity (Stringer et al., 2019).

At the mesoscale level, there are reported similarities in the characteristics of evoked and spontaneous activities. For example, Arieli et al., 1995 stated that the amplitude of the ongoing spontaneous activity was similar to the one in evoked responses in a portion of the anesthetized cat visual cortex (Arieli et al., 1995). Moreover, Kenet et al., 2003 suggested that spontaneous activity patterns are similar to the orientation maps observed in visually evoked responses in anesthetized cats (Kenet et al., 2003). More recently, it has been shown that these maps emerge only in anaesthetized states but not in awake states (Omer et al., 2019). Similarly, Han et al., 2008 showed that the propagation patterns of spontaneous activity resembles recent sensory-evoked patterns when measured in a portion of the visual cortex in anesthetized rats (Han et al., 2008). At the macro-scale level, the fluctuations of the BOLD signal in functional magnetic resonance imaging (fMRI) in humans has been reported to be larger during spontaneous activity (resting state) than during task-driven activity (He, 2011; Ponce-Alvarez et al., 2015; Ito et al., 2019).

Some of the contradictory results on the relationship between evoked and spontaneous activity in previous studies can be explained by methodological differences. For example, these studies used different animal models, which can affect certain characteristics of cortical processing (e.g. theta oscillations in humans are significantly slower than in rodents) (Alloway et al., 1993; Jacobs, 2014). Moreover, the recording techniques used in these studies have important intrinsic differences in spatial and temporal resolutions, which in turn might impact the measurements to compare evoked and spontaneous activity (Menon and Kim, 1999). Furthermore, there is also wide diversity in the preparations used in these studies, such as brain slices or *in vivo* recordings, which have distinct activity dynamics and might impact the conclusions (Azzarelli et al., 2017). Also, the brain states in which the recordings were performed might be different. For example, the cortical dynamics of anesthetized or sleeping animals have remarkable differences to the ones from awake or desynchronized brain states (Sellers et al., 2015). Finally, the sensory modalities from which the brain activity was recorded from might not be directly comparable. For example, it is known that locomotion can modulate evoked responses in auditory and visual cortices in opposite directions (Niell and Stryker, 2010; McGinley et al., 2015; Yang et al., 2020).

In this study, we used VSD and glutamate mesoscale wide-field imaging of most of the right cerebral hemisphere in lightly anesthetized and awake mice, respectively to evaluate whether the complexity of cortical activity patterns was different during spontaneous and evoked activity. Optical flow and fractal dimension analyses were used to characterize and compare the spatiotemporal dynamics of sensory-evoked activity to to motifs of spontaneous activity that originate in the same cortical region using evoked activity-derived template matching (Mohajerani et al., 2013; Afrashteh et al., 2017). Overall, this technique offers the possibility of monitoring activity over large portions of the cortical mantle at high spatiotemporal resolution which, in turn, permits the application of methods to characterize the cortical activity dynamics in both regimes. In contrast with previous studies, we used different sensory modalities (tactile, auditory, and visual), several stimulation strengths, and brain states, which provide a more complete comparison of ongoing and evoked cortical activity.

We found that the response amplitude, propagation speed, and extent of propagation of the evoked activity has a positive correlation with the stimulus intensity. The speed, direction stability, and extent of propagation of spontaneous activity was also positively correlated with the amplitude. Our results demonstrate that, for the above-mentioned parameters, sensory-evoked responses at low stimulus strengths are similar to motifs of spontaneous activity and that this similarity diminishes as stimulus strength increases. Motifs were defined as spatiotemporal patterns within spontaneous activity that matched spatiotemporal templates constructed from sensory evoked responses. Finally, we showed that the repertoire of directions and trajectories of propagation of sensory-evoked activity is smaller than the one from spontaneous activity.

## 2 Materials and Methods

### 2.1 Animals and Surgery

Animal housing, surgery, and all experimental protocols were approved by the University of Lethbridge Animal Care Committee in line with the guidelines from the Canadian Council for Animal Care. For anesthetized preparations, adult male and female C57BL/6J mice (n=5 animals for forelimb and hindlimb stimulation experiments and n=4 animals for visual stimulation experiments) around ∼3 months of age and weighing approximately 25g were used. Mice were anesthetized with isoflurane (1.0–1.5%) for induction and during surgery, and a reduced concentration during data collection (0.5–1.0%). A 7 × 6 mm unilateral craniotomy (bregma +2.5 to -4.5 mm, lateral to the midline 0 to 6 mm) was made, and the dura was removed, as described previously (Kyweriga and Mohajerani, 2016). The body temperature was maintained at 37°C during surgeries and subsequent imaging sessions. For awake preparations, n=5 adult mice (2 males and 3 females) Ai85-CamKtTA-EMX1-Cre (cross of B6; 129S-Igs7tm85.1 (tetO-gltI/GFP*) Hze/J and B6.129S2-Emx1tm1(cre)Krj/J) with 5–8 months of age and weighing approximately 26g were used. These mice express intensity-based glutamate sensing fluorescent reporter (iGluSnFR) in excitatory neurons and glial cells of cortex (Xie et al., 2016; Karimi Abadchi et al., 2020). One week prior to performing wide-field glutamate imaging, a surgery was done to expose the cranial bone and for implanting a head-plate. A large portion of the cranial bone was exposed by removing the skin. The cranial bone was then cleaned and metabond was applied on top. Finally, a glass coverslip was placed on top to keep the surface clear from accumulating debris from the animal’s home cage and the environment.

### 2.2 Data acquisition

#### 2.2.1 Wide-field optical imaging

Wide-field optical imaging of summated cortical synaptic or voltage activity was utilized to capture the mesoscale dynamics of the cortex in both anesthetized and awake preparations. For voltage-sensitive dye imaging in anesthetized mice, the dye RH-1691 (Optical Imaging, New York, NY) (Shoham et al., 1999), was dissolved in HEPES-buffered saline solution (0.5 mg/ml) and applied to the exposed cortex for 30-40 min, as described previously (Mohajerani et al., 2010; Mohajerani et al., 2011; Lim et al., 2012; Lim et al., 2013). The unabsorbed dye was carefully washed out and the brain was covered with 1.5% agarose in HEPES-buffered saline solution and sealed with a glass coverslip. VSD was excited using a red LED (627 nm center, Luxeon Star LEDs Quadica Developments Inc., Alberta, Canada) and excitation filter (630±15 nm, Semrock, New York, NY). The VSD signal was passed through an emission filter (688±15 nm, Semrock, New York, NY). Images were taken through a macroscope composed of front-to-front pair of video lenses (8.6×8.6 mm field of view, 67 µm per pixel). The focal plane of the camera was 0.5– 1 mm below the cortical surface. The images were captured by a 12-bit charge-coupled device (CCD) camera (1M60 Pantera Dalsa, Waterloo, ON) and an EPIX E8 frame grabber with XCAP 3.8 imaging software (EPIX, Inc., Buffalo Grove, IL) at 150 Hz frame rate. For glutamate imaging, iGluSnFR was excited using a blue LED (470nm center, Luxeon Star LEDs Quadica Developments Inc., Alberta, Canada) filtered through an excitation filter (470±20nm, Semrock, New York, NY). The fluorescent signal emitted from cortical activity was collected after passing through an emission filter (542±27nm). Images were taken with the same macroscope camera as described above.

#### 2.2.2 Recording of spontaneous and evoked activity

For anesthetized preparations, spontaneous activity was recorded for at least 20 mins prior to recording evoked activity as sensory stimulation could alter the characteristics of spontaneous activity (Han et al., 2008). To electrically stimulate left forelimb and hindlimb paws, a thin needle (0.14 mm) was inserted into the ventral surface of each paw. Different levels of current were presented in ascending order to stimulate the paws for 1ms duration (0.01 to 1.5mA; Supplementary Fig. SM1A). The stimulation intensities were categorized into three classes: low (Lo), medium (Med), and high (Hi) strength as follows: the low and high stimulation levels were identified as the minimum levels that evoke a detectable VSD response and a saturated response, respectively. The medium stimulation level was a level between low and high stimulation levels that generated a response with clear distinction from low and high responses (Supplementary Fig. SM1A). A 1-ms pulse of combined blue and green light was used as visual stimulation. For fore and hind paw stimulation, 10–20 trials of each stimulation intensity level were given with an interstimulus interval of 10s. For visual stimulation experiments, VSD response distributions were calculated from 100–200 trials of stimulation with an interstimulus interval of 10s.

For awake preparations, spontaneous activity was recorded similarly as described above. Evoked activity was recorded by presenting a tone of 12 kHz frequency for a duration of 50ms. The sound levels for Lo and Hi stimulation levels were 40 and 80 dBSPL respectively. 10-20 trials of stimulation were presented with an intertrial interval of 10-12s. Tones were generated using a Tucker-Davis Technologies Inc. (TDT, Alachua, FL, USA) RX6 processor and delivered to animal’s left ear via a free-field ES1 (TDT Inc.) speaker.

### 2.3 Data analysis

#### 2.3.1 Data preprocessing

All preprocessing was done in Matlab® (Mathworks, Natick, MA, USA). To do fair comparisons between evoked responses and spontaneous activity, all recordings (image frames) were treated using similar preprocessing procedures. As staining of non-flat cortical surface with voltage-sensitive dye is likely non-uniform, a regional bias in the captured signal is possible. To overcome this bias, the signal was normalized to a baseline (ΔF/F_0_). The baseline was estimated for both, no-stimulation trials and spontaneous activity, using the “locdetrend” function from the Chronux toolbox (Bokil et al., 2010). The ΔF/F_0_ time series of the VSD signal was then filtered with a zero-phase lag Gaussian low-pass filter (6 dB attenuation at 25 Hz frequency) while the glutamate signal was filtered with 6 Hz low-pass filter to eliminate the effects of hemodynamics that appears at 8-14 Hz. At the end, a two-dimensional spatial Gaussian filter (σ ≈ 67 *μm*) was applied. Each evoked response trial was preprocessed separately and then averaged to get the average response of the corresponding cortical activation and stimulation level.

#### 2.3.2 Optical flow analysis

To capture the spatiotemporal dynamics of VSD and glutamate recordings, we used the Combined Local-Global (CLG) method implemented in Optical-Flow Analysis Toolbox (Afrashteh et al., 2017). Using this method, velocity vector fields were estimated for each time point. For each pixel in a frame, we calculated a velocity vector that depicts the instantaneous speed and direction of activity propagation.

#### 2.3.3 Identification of spontaneous activity motifs

For anesthetized mice, spontaneous activity patterns were compared to patterns of evoked activity in the primary forelimb somatosensory (FLS1), hindlimb somatosensory (HLS1), and visual cortices (VC) while for awake mice, similar comparisons were done for activity in the auditory cortex (AC). Motifs of spontaneous activity were identified using the template matching method used in (Mohajerani et al., 2013). Briefly, for each sensory modality the template was defined as the set of three frames after the onset of the evoked activity elicited by Hi stimulus level (Supplementary Fig. SM1B). Then, we calculated the Pearson correlation coefficients (PCC) between these templates and spontaneous activity (Fig. SM1C and D). Frames of spontaneous activity with a PCC greater than a given threshold were considered to be a ‘match’ to the evoked templates. To determine the threshold value, the maximum of PCC between all sensory templates (hindlimb, forelimb, visual, auditory and whisker) were measured, and the results were multiplied by a constant factor (1.34) (Mohajerani et al., 2013). The consecutive events with PCC peaks above given threshold were also screened to be separated by at least 100msec. Frames around these time points were then selected as spontaneous activity events or motifs. To determine the onset of a spontaneous activity motif, local minima were found within 50 frames around the PCC peak and fitted with a cubic polynomial curve (as shown in supplementary Fig. SM2A). The onset of a spontaneous activity motif was considered one frame prior to the intersection of the fitted curve and the original signal. We validated this methodology by estimating stimulus onsets for all trials of evoked activity and comparing with actual values. Most of the differences between estimated and actual onsets were close to 0 (supplementary Fig. SM2B, SM2C, SM2D). The estimation was less accurate for glutamate imaging (supplementary Fig SM2E). For all subsequent analyses, “estimated” onsets for both evoked and spontaneous activity were used. To determine the trajectory of the activity we needed to determine the end of such activity. For this, we considered the activity peak as end point since after this frame the activity starts to diminish. Note that for finding the spontaneous motifs in the forelimb and hindlimb regions, the threshold value for finding PCC peaks (see above) was defined individually for each animal as the mean PCC between forelimb and hindlimb-evoked activity (first three frames) for all stimulus intensities (total 9 values). If the mean PCC value was less than 0.4, 0.4 threshold value was used.

#### 2.3.4 Measurement of amplitude, time-to-peak, and propagation speed and direction

To determine characteristics of evoked and spontaneous activity such as amplitude, and time-to-peak, speed and direction in a cortical area (e.g. FLS1, HLS1, VC, or AC), a region-of-interest (ROI) was first defined and time series of ΔF/F_0_ for all pixels within the ROI were averaged to obtain one ΔF/F_0_ time series which was used in subsequent analyses. A ROI was defined as the area for which ΔF/F_0_ values are above the mean+SD of all ΔF/F_0_ values in that frame. The center of the PTA ROI was located anatomically as -2mm anterior-posterior and +1.5mm lateral from the bregma and with the size of 0.5×1mm^2^. To calculate the amplitude of sensory-evoked responses, the mean of the baseline was subtracted from the maximum of ΔF/F_0_ values within the first 10 frames (67ms for VSD and 100ms for glutamate experiments) after stimulus onset. Similarly, to calculate the amplitude of spontaneous activity motifs, the mean of baseline activity was subtracted from the peak ΔF/F_0_ within ±15 frames around the peak PCC used for identifying a motif (see above). For both evoked and spontaneous activity, the baseline was considered as ΔF/F_0_ of 10 frames before the onset of activity. The time-to-peak was calculated as the time of the peak minus the time of the estimated activity onset.

To calculate the propagation speed and direction of the activity from the optical flow analysis, we averaged the velocity vectors of all pixels in each ROI for each time frame. Speed and direction of propagation were then determined from the velocity vector with the largest magnitude after the estimated activity onset and within ±15 frames around the peak of ΔF/F_0_ signal. For determining how stable the directions of propagation were for a spontaneous activity motif, we defined the propagation direction stability (Figures 7 and S7) and calculated as follows. First, for each frame, we calculated the cosine of the angles between the average velocity vector at peak speed and the average velocity vectors in the frames before and after such frame (±6 and ±10 frames around the peak speed frame for VSD and glutamate imaging respectively). Then, we used the average cosine value to estimate how similar the directions of propagation of the motif to its direction at peak speed were. Therefore, the propagation direction stability measured this way has values between - 1 and 1 with values closer to 1 indicating that on average, the direction of propagation of spontaneous activity motif was similar to its direction at peak speed.

#### 2.3.5 Determination of propagation trajectories/paths of activity

To compare the propagation of sensory-evoked and spontaneous activity motifs, trajectories were calculated starting from the onsets of activity to temporal locations of their peak amplitude (Fig. SM2F). To estimate the activity trajectory, we first determined the extent of activity (i.e. onset and offset) and its centroid for each frame using contours identified by ΔF/F_0_ values above mean + m×SD of all ΔF/F_0_ values in that frame. The value of “m” was 0.6 and 0.5 for the first and subsequent frames respectively. Next, one spatial point was selected in each frame to represent the “location of activity” in that frame. The trajectory of the activity propagation was then defined as the sequence of such activity locations across frames. In the first frame, the centroid of the activity contour was selected as the “location of activity”. For all subsequent frames, the location of activity was defined as the intersection of the contour line and the extrapolated directional vector joining the centroid of activity in the previous frame to centroid of activity in the current frame (Fig. SM2F).

### 2.4 Statistical Analysis

#### 2.4.1 Statistical Tests

All statistical analyses were done in Matlab®. Paired sample t-test, Wilcoxon rank sum test, or repeated measures ANOVA with Greenhouse-Geisser adjustment for the potential lack of sphericity and post-hoc Tukey-Kramer correction for multiple comparisons were used. We report F-values, p-values and the estimates of the effect size (partial η^2^). Also, in Figures 7 and S7 we fitted a linear regression model to each of the scatter plots. The linear model coefficients and their p-values are reported. The type of test used is indicated in the text as well as in figure captions. Error bars represent the standard error of mean (SEM). *, **, and *** indicate significance levels of *p* < 0.05, *p* < 0.01, and *p* < 0.001 respectively.

#### 2.4.2 Bootstrap Method to generate Distributions of Hausdorff Fractal Dimension

The Hausdorff fractal dimension measures the roughness (complexity) of a trajectory path. Trajectories that occupy larger space exhibit higher fractal dimension (Singh et al., 2016). We determined the Hausdorff fractal dimensions using the box counting method for each trajectory heat map which encode the percentage of the number of activity that passed through a certain point on the cortical surface. The heat maps were first binarized using Otsu’s method (Otsu, 1979) before determining fractal dimensions. For each animal and stimulus type (e.g. forelimb stimulation), we generated a distribution of Hausdorff fractal dimension using bootstrapping over trials from all levels of stimulus strength. Once we calculated these distributions over all trials from all stimulus levels, we calculated the trajectory for each stimulation trial. At each iteration of this bootstrapping method, we randomly selected the corresponding trajectories of 10 trials and their average trajectory was used to represent the trajectory in this condition. Then, we calculated the fractal dimension for the average trajectory. This process was done 500 times and we estimated the probability density function (PDF) of the calculated fractal dimension. Analogously, we used the same procedure to estimate the PDF of spontaneous motifs for each animal and stimulus-type motifs.

## 3 Results

### 3.1 Response amplitudes of sensory-evoked cortical activity are larger than the amplitudes of motifs of spontaneous cortical activity in both anesthetized and awake mice

We investigated how changes in stimulus strength affect the amplitude of sensory-evoked activity and compared it with amplitudes of spontaneous activity events or motifs (see Methods). In previous studies (Petersen et al., 2003b; MacLean et al., 2005; Han et al., 2008; Luczak et al., 2009), the amplitude of spontaneous activity was found to be comparable to the amplitude of evoked responses. Works on the topographic organization and processing characteristics of different sensory cortices using fMRI, electrophysiology, and optical imaging (Jancke et al., 1998; Shoham et al., 1999; Spenger et al., 2000; Petersen et al., 2003a; Ferezou et al., 2007; Polley et al., 2007; Guo et al., 2012), suggested that increasing the stimulus intensity induces a larger response amplitude at the neuronal network level. Here, we revisited the study of the the relation between the amplitude of spontaneous motifs and evoked activity elicited by multiple stimulus strengths to clarify this discrepancy in both, anesthetized and awake animals.

We used voltage-sensitive dye (VSD) imaging in anesthetized mice and modeled different levels of somatosensory stimulation by injecting several different electrical current levels, low (Lo), medium (Med) and high (Hi), to the fore and hind paws. Different level of stimuli strength used here may mimic the statistics of the sensory input experienced by animals during natural behaviour (Simoncelli, 2003; Carriot et al., 2017). Anaesthesia provided a long-lasting, stable baseline brain state that was free of active behaviours and voluntary mental processes that may contaminate the analysis of purely spontaneous activity from which to make our observations (Musall et al., 2019; Stringer et al., 2019). We measured evoked cortical responses in the contralateral cerebral hemisphere for multiple trials of forelimb or hindlimb stimulation. We also recorded spontaneous activity and identified events which were similar to patterns of cortical activation elicited by forelimb and hindlimb stimulation. These spontaneously occurring motifs were determined using a template matching method with the templates extracted from evoked responses elicited by Hi forelimb and hindlimb stimulus strength (see Methods & Supplemental Fig. SM1 B-D). Using this approach, we identified the spontaneous activity motifs originated in the primary fore or hindlimb somatosensory (FLS1 or HLS1) cortices (top rows in Fig. 1A and S1A) and propagated following different trajectories on the cortical surface. Sensory-evoked activity increased in amplitude and spread when stimulus intensities were increased (Fig. 1A and supplementary Fig. S1A). For low level forelimb and hindlimb stimuli, the response was local to the forelimb and hindlimb areas of the primary and secondary somatosensory cortices respectively, while for medium and high stimuli, the spatial extent of the evoked response was larger than the primary forelimb and hindlimb somatosensory areas. Moreover, for forelimb stimuli, after an initial expansion of the evoked response in the FLS1 area, activity traveled in the medio-caudal direction passing through the HLS1 area (Fig 1A). Similarly, the spread of the initial hindlimb-evoked response increased with stronger stimuli (supplementary Fig. S1A).

**Fig. 1.**
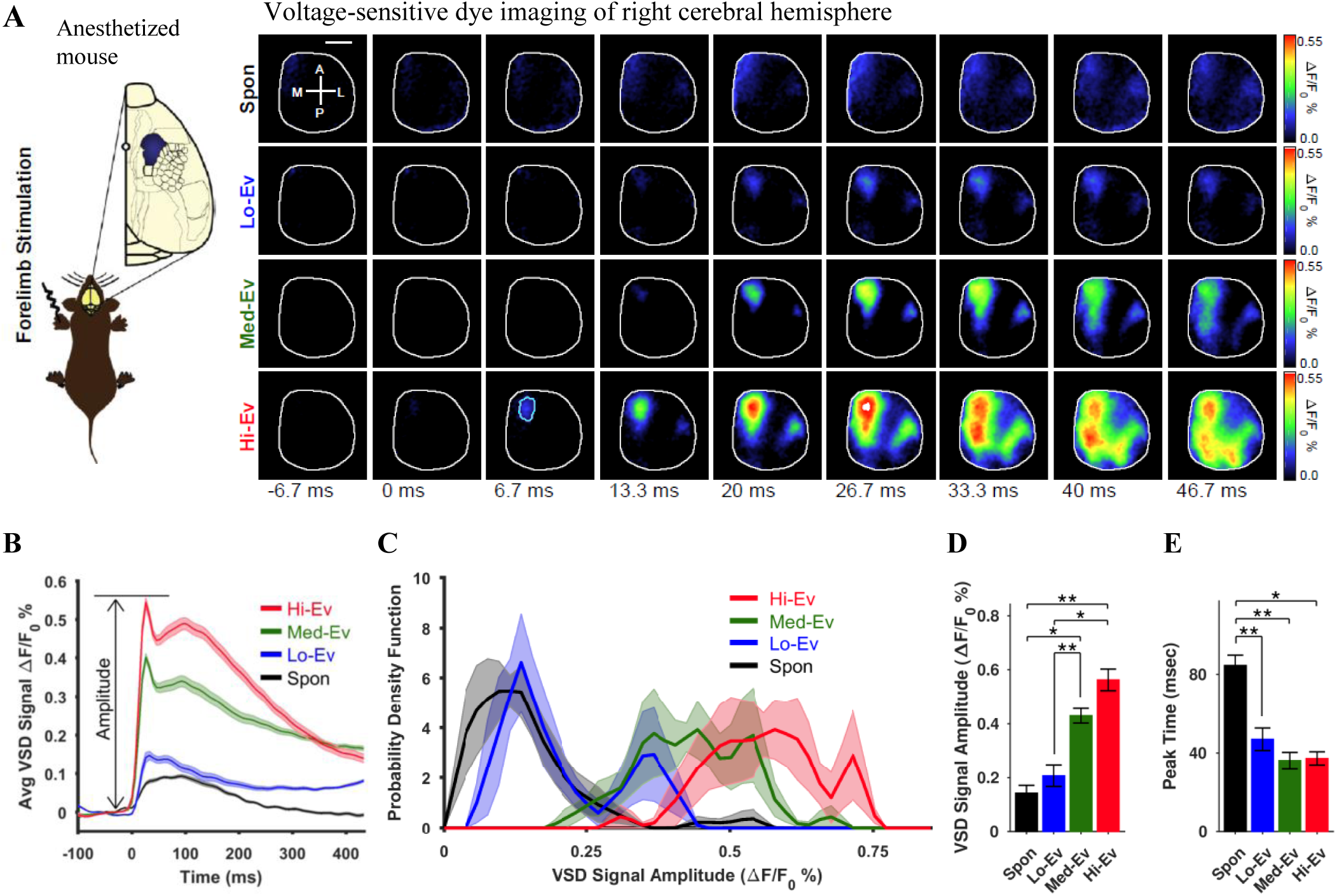
Amplitude of forelimb-evoked activity is larger than that of spontaneous activity in anesthetized mice. **(A)** Experimental paradigm (left) and montages of VSD imaging (right). Montage top row shows a representative average of forelimb-motifs in spontaneous (Spon) activity (averaged from 10 motifs). Bottom three rows show average evoked (Ev) cortical activity in response to forelimb stimulation with low (Lo), medium (Med), and high (Hi) stimulus strengths (averaged over 10 trials). The primary forelimb region of interest (FLS1-ROI) is outlined in the third column of the bottom row. Compass lines indicate anterior (A), posterior (P), medial (M) and lateral (L) directions. Scale bar is 1 mm. **(B)** Plots of the average VSD signal in the FLS1-ROI across animals (n=5). The lines represent the mean over animals calculated from the averages over trials for individual animals. Thick lines indicate the mean values while shaded regions indicate SEM across animals. **(C)** Average distributions of VSD signal amplitudes in the FLS1-ROI (normalized to the percentage of occurrences). VSD Signal Amplitude = (peak ΔF/F_0_ – (mean of baseline) as depicted in (**B**). Shaded regions denote the SEM across animals. **(D**-**E)** Mean±SEM values of VSD signal peak amplitudes and time-to-peak (time from the onset to the peak of response) respectively. * and ** indicate p<0.05 and p<0.01 respectively repeated measure ANOVA with post-hoc Tukey-Kramer correction.

For a quantitative analysis, we first determined the average value of the VSD signal over time in an ROI (FLS1 or HLS1) by finding the mean ΔF/F_0_ for all pixels within an ROI at each time point. In this way, time series of VSD signals for an ROI were determined for all trials of evoked activity and for all motifs of spontaneous activity. The average over trials (evoked-activity) and motifs (spontaneous activity) for all animals (n = 5) of these VSD signals is shown in Fig. 1B for FLS1 ROI (supplementary Fig. S1B for hindlimb stimulation and HLS1 ROI). It is quite evident that motifs of spontaneous activity have the lowest ΔF/F_0_ values while these values increase with increasing stimulus levels for evoked activity. Next, we calculated the amplitudes of these signals by subtracting the mean value over the baseline from the peak ΔF/F_0_ (see Fig. 1B) and observed that the amplitudes of evoked activity increased with the intensity of the stimulus (Fig. 1B-C, and S1B-C). Note that the mean amplitudes of spontaneous activity motifs were similar to those of evoked activity elicited with Lo stimulus levels while they were significantly smaller than amplitudes of evoked activity elicited with Med and Hi stimulus levels (Fig. 1D for forelimb, repeated measures ANOVA: F-value = 29.075921, p = 0.0000086849, η^2^ = 0.879066; post-hoc: spontaneous vs Med p = 0.018772, spontaneous vs Hi p = 0.0059376; Fig. S1D for hindlimb, repeated measures ANOVA: F-value = 27.93219, p = 0.0000107044, η^2^ = 0.874735; post-hoc: spontaneous vs Med p = 0.048948, spontaneous vs Hi p = 0.011894). We also quantified the time-to-peak (time from stimulus onset to the VSD signal maximum) and found that spontaneous activity motifs were the slowest to reach their peak levels as compared to responses of evoked activity (Fig. 1E for forelimb, repeated measures ANOVA: F-value = 20.520079, p = 0.0000512563, η^2^ = 0.836868; post-hoc: spontaneous vs Lo p = 0.0051397, spontaneous vs Med p = 0.0067666, spontaneous vs Hi p = 0.011896; Fig. S1E for hindlimb, repeated measures ANOVA: F-value = 8.992695, p = 0.0021416665, η^2^ = 0.692135; post-hoc: spontaneous vs Hi p = 0.038979). In a separate cohort of anesthetized mice (n = 4), we also compared the amplitudes and time-to-peak of visual evoked activity (elicited with only one stimulus level) with those of visual motifs in spontaneous activity (supplementary Fig. S2). The mean amplitude of evoked activity was significantly larger than that of spontaneous activity events (Fig. S2D, paired sample t-test p = 0.012257). The time-to-peak for evoked activity was significantly smaller than that for visual motifs in spontaneous activity (Fig. S2E, paired sample t-test p = 0.0078674).

To compare the amplitudes and temporal dynamics of sensory-evoked and spontaneous activity in the awake brain state, we used glutamate (Glu) imaging and auditory stimulation with two levels of sound volume (Lo and Hi). The preprocessing to find Glu ΔF/F_0_ values was done in a similar fashion as for VSD imaging except that Glu signal was filtered using a bandpass filter (see methods). Spontaneous activity motifs were also determined in a similar fashion as described above for VSD imaging. The auditory-evoked response originated in the primary auditory cortical region (AC) and expanded into other cortical regions (montage Fig. 2A). The Glu signal was the largest for evoked response elicited by Hi stimulus level. The average Glu signal (Fig. 2B; average over ROI pixels, trials, and animals) was also the largest for evoked activity elicited by Hi stimulus level. However, average Glu signals for evoked activity elicited by Lo stimulus level was comparable to average Glu signal for spontaneous activity motifs. Next, we compared the amplitudes of the Glu signal in a similar way as for VSD signal. The amplitudes of auditory motifs in spontaneous activity were significantly smaller than those of evoked activity elicited with Hi stimulus levels (Fig. 2C-D, repeated measures ANOVA: F-value = 19.906767, p = 0.0007837121, η^2^ = 0.832683; post-hoc: spontaneous vs Hi p = 0.0040670) but were similar to those elicited by Lo stimulus level (p = 0.93913). The time-to-peak for motifs of spontaneous activity was significantly larger than the time-to-peak for responses of evoked activity with Hi and Lo auditory stimulus levels (Fig. 2E, repeated measures ANOVA: F-value = 201.184997, p = 0.0000001444, η^2^ = 0.980505; post-hoc: spontaneous vs Lo p = 0.00024736, spontaneous vs Hi p = 0.00030659). These results, from both anesthetized and awake preparations, thus suggest that the amplitudes of spontaneous activity motifs are smaller than those of evoked activity and their temporal dynamics is slower.

**Fig. 2.**
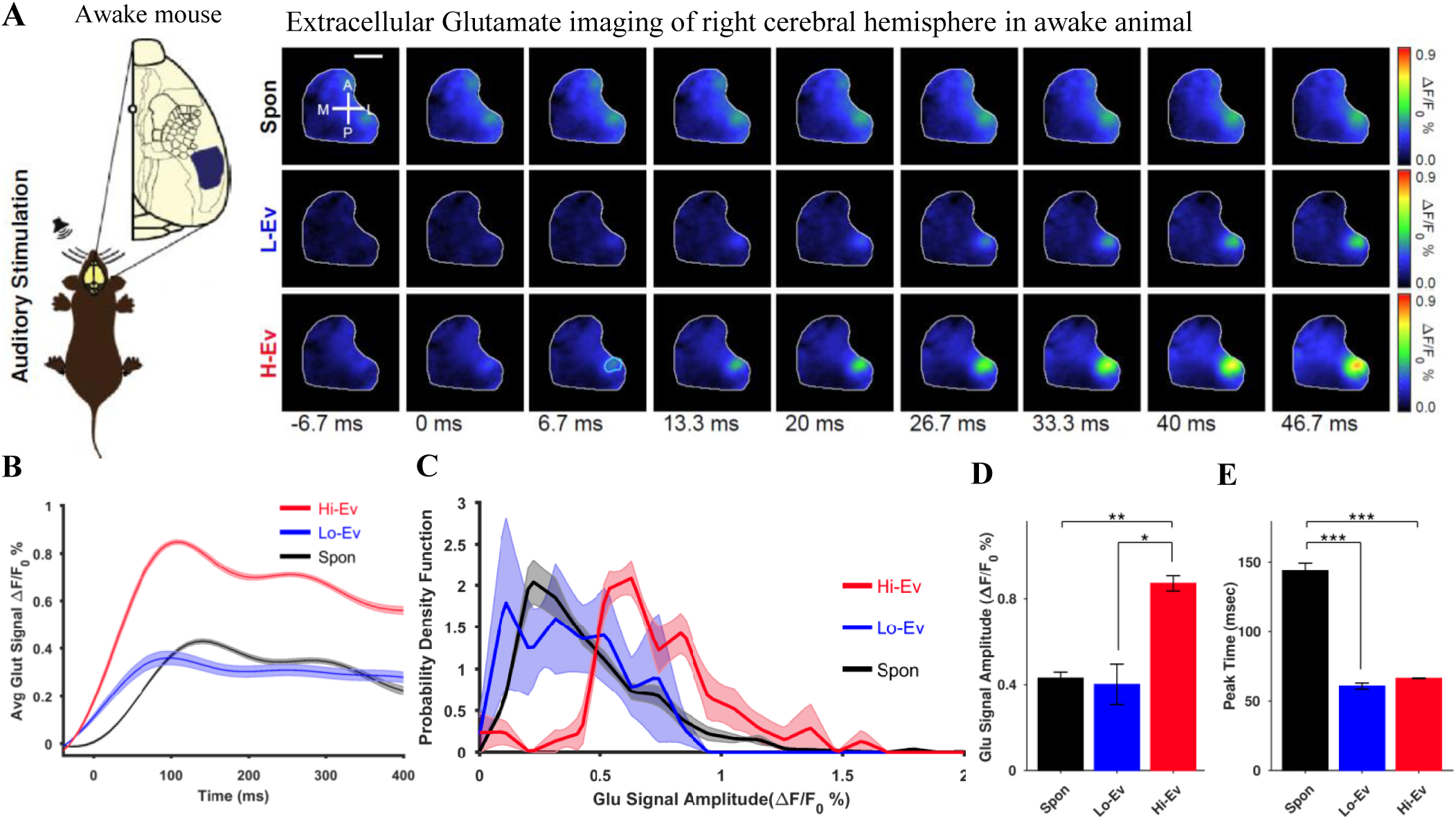
Amplitude of tone-evoked activity is larger than that of spontaneous activity in awake mice. **(A)** Experimental paradigm (left) and montages of extracellular glutamate imaging (right). Montage top row shows a representative average motif of auditory spontaneous (Spon) activity motifs. Bottom two rows show average evoked (Ev) cortical activity in response to auditory stimulation with low (Lo) and high (Hi) stimuli strengths. The primary auditory region of interest (AC-ROI) is outlined in the third column of the bottom row. Compass lines indicate anterior (A), posterior (P), medial (M) and lateral (L) directions. Scale bar is 1 mm. **(B)** Plots of the average glutamate signal in the AC-ROI (n=5 animals). The plots represent average over animals calculated from averages over trials for individual animals. Thick lines indicate the mean values while shaded regions indicate SEM across animals. **(C)** Average distributions of glutamate signal amplitudes in the AC-ROI (normalized to the percentage of occurrences). Glutamate signal amplitude = (peak ΔF/F_0_) – (mean of baseline). Shaded regions represent the SEM. **(D-E)** Bar graphs show the mean±SEM values of glutamate signal amplitude peaks and time-to-peak respectively. *, **, and *** indicate p<0.05, p<0.01, and p<0.001 respectively for a repeated measure ANOVA with post-hoc Tukey-Kramer correction.

### 3.2 The propagation speed of sensory-evoked cortical activity is larger than that of motifs of spontaneous cortical activity in anesthetized and awake mice

We next investigated how stimuli of different strengths affects the speed of propagation of evoked activity waves on the cortical surface and whether the speeds of sensory-evoked activity are larger than those of spontaneous activity. Given that, behaviorally, animals must attend to salient stimuli for survival (Corbetta et al., 2008), we hypothesized that the propagation speeds of sensory-evoked activity would increase with highly salient stimuli and will be higher than the speeds of spontaneous activity. To address these questions, we quantified the propagation speed of sensory-evoked activity for forelimb and hindlimb modalities in anesthetized mice with VSD imaging. Similarly, we quantified the speed of auditory evoked responses in awake mice with glutamate imaging. To determine velocity vector fields on VSD and glutamate imaging recordings, we applied optical flow analysis (Afrashteh et al., 2017) to obtain the instantaneous speed and direction of propagation (Fig. 3A).

**Fig. 3.**
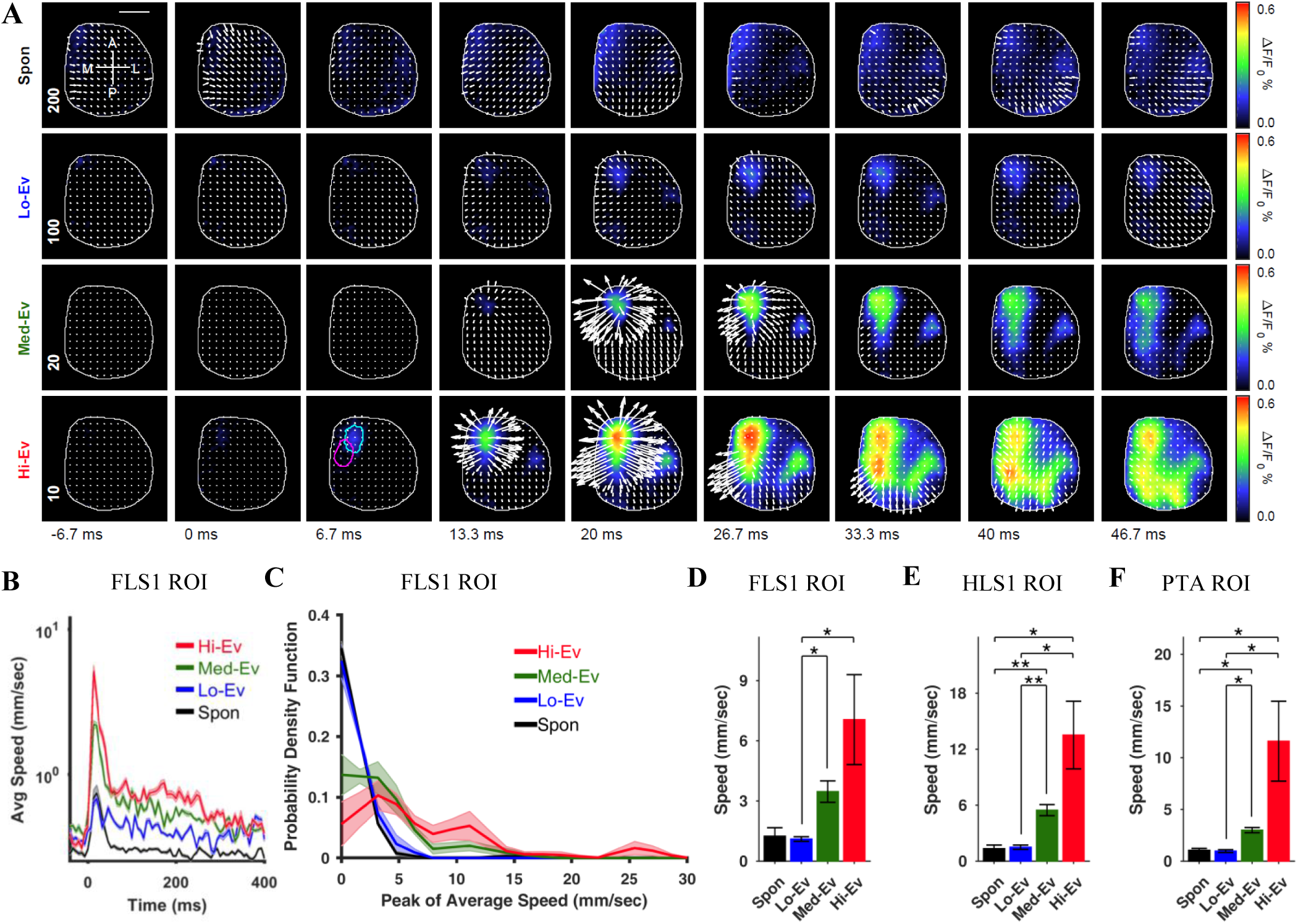
Propagation speed of forelimb-evoked activity is larger than that of spontaneous activity in anesthetized mice. **(A)** Montage of representative examples of motifs of forelimb spontaneous (Spon) activity (top row) with overlaid velocity vector fields determined by optical flow analysis. The bottom three rows show similar montages for average evoked (Ev) activity in response to contralateral forelimb stimulation with low (Lo), medium (Med), and high (Hi) stimuli strengths. Primary sensory forelimb (FLS1) and hindlimb (HLS1) ROIs are outlined in the third column of the bottom row. Compass lines indicate anterior (A), posterior (P), medial (M) and lateral (L) directions. Scale is 1 mm. Numbers in the first column indicate scale factor for drawing velocity vector fields. **(B)** Plots of the average speed signal in the FLS1-ROI (n=5 animals). Note that the scale in the y axis is logarithmic. Shaded regions indicate SEM across animals. **(C)** Average distributions of peak average speeds in the FLS1-ROI (normalized to the percentage of occurrences). Shaded regions represent the SEM across animals. **(D-E)**, and **(F)** Mean±SEM values of peak from average speeds in the FLS1, HLS1, and PTA ROIs respectively. Note that different y-axis scales are used for each ROI. * and ** indicate p<0.05 and p<0.01 respectively for a repeated measure ANOVA test with post-hoc Tukey-Kramer correction.

For anesthetized mice and somatosensory modalities (forelimb and hindlimb), the instantaneous speeds increased with stronger stimuli (Fig. 3A and S3A). Instantaneous speeds for forelimb motifs in spontaneous activity and for responses elicited by Lo stimulus strength, however, were very small (notice different scale factors used for plotting velocity vector fields in first column in Fig. 3A and S3A). Furthermore, we calculated the average speed over time by finding the magnitude of the vector sum of the velocity vectors of all pixels within an ROI (e.g. FLS1 or HLS1) for each time point calculated over the average response of all trials. The average over animals (n = 5) was then determined and shown in Fig. 3B for FLS1 ROI (and in supplementary Fig. S3B for hindlimb stimulation and HLS1 ROI). The average speeds were the lowest for spontaneous activity motifs and increased with increasing stimulus levels for evoked activity. The peak speeds in the corresponding ROIs were also determined from the time series of average speeds. The distributions of these peak speeds are shown in Fig. 3C (and in supplementary Fig. S3C for hindlimb stimulation). Motifs of spontaneous activity had peak speeds similar to those of responses of evoked activity elicited with Lo stimulus strength. The speeds increased with increasing stimulus levels (Fig. 3C and 3D) in the FLS1 ROI. The mean peak speed of evoked activity elicited by Hi and Med stimulus strength was significantly larger than the mean peak speed of evoked activity elicited by Lo stimulus strength (Fig. 3D, repeated measures ANOVA: F-value = 9.226881, p = 0.0019301139, η^2^ = 0.697586; post-hoc: Med vs Lo p = 0.012693, Hi vs Lo p = 0.036275). Since forelimb evoked activity elicited by Med and Hi stimulus levels originated in the FLS1 ROI but traveled in the medio-caudal direction on the cortical surface, we also compared propagation speeds in the HLS1 and posterior tegmental area (PTA) (posterior parietal cortex) ROIs. The difference in speeds of sensory-evoked with Med and Hi stimulus strengths and spontaneous activity was significant in the HLS1 and PTA ROIs (Fig. 3E for HLS1, repeated measures ANOVA: F-value = 19.027242, p = 0.0000743228, η^2^ = 0.826293; post-hoc: Med vs spontaneous p = 0.0068222, Hi vs spontaneous p = 0.036175; 3F for PTA, repeated measures ANOVA: F-value = 14.780107, p = 0.0159957714, η^2^ = 0.787009; post-hoc: Med vs spontaneous p = 0.041526, Hi vs spontaneous p = 0.038762). Also, for HLS1 and PTA ROIs the mean speed for Med and Hi strength of sensory forelimb stimulation is significantly higher than the mean speed of propagation for Lo stimulus level (Fig. 3E for HLS1, Med vs Lo p = 0.0062288, Hi vs Lo p = 0.018329; 3F for PTA, Med vs Lo p = 0.017250, Hi vs Lo p = 0.036929). For hindlimb stimulation also similar trend is observed in which Hi stimulus level evokes significantly larger speed compared to Lo and Med stimulus strength and spontaneous activity in all three ROIs (Fig. S3D for HLS1, repeated measures ANOVA: F-value = 28.555109, p = 0.0008278413, η^2^ = 0.877131; post-hoc: Hi vs Lo p = 0.0067064, Hi vs Med p = 0.020179, Hi vs spontaneous p = 0.017031; Fig. S3E for FLS1, repeated measures ANOVA: F-value = 23.909933, p = 0.0000238001, η^2^ = 0.856682; post-hoc: Hi vs Lo p = 0.016317, Hi vs Med p = 0.0078756, Hi vs spontaneous p = 0.025145; Fig. S3F for PTA, repeated measures ANOVA: F-value = 17.738107, p = 0.0069307567, η^2^ = 0.815991; post-hoc: Hi vs Lo p = 0.026711, Hi vs Med p = 0.027323, Hi vs spontaneous p = 0.044441). We also performed optical flow analysis for visual cortical stimulation in a separate cohort of anesthetized mice (n = 4) and found that peak speeds of evoked activity are larger than those of spontaneous activity motifs (Fig. S4, paired sample t-test p = 0.032798). These results thus suggest that in anesthetized mice, the propagation speeds for spontaneous activity are comparable to those in evoked activity elicited with Lo stimulus levels but are smaller than those of evoked activity elicited with Hi stimulus levels.

Next, we compared the propagation speeds of spontaneous activity with those of sensory evoked activity in awake mice. The velocity fields for spontaneous and evoked activity at Lo stimulus strength had vectors with comparable magnitudes while sensory evoked propagations with Hi stimulus level showed velocity vectors with larger magnitudes (Fig. 4A, see scale factors for plotting vector fields and compare with scales in Fig. 3A). The average speeds (over trials and animals) were determined as described above and shown in Fig. 4B. All the graphs show a baseline speed close to zero and then a rise in speed after the stimulus onset or spontaneous motif onset which gets to its peak and then declines towards baseline. The distribution of peak speeds determined from the average speed curves (Fig. 4C) were similar for evoked responses by Lo stimulus level and spontaneous motifs with no significant difference between means of peak speeds in AC ROI (Fig. 4D, repeated measures ANOVA: F-value = 37.011210, p = 0.0000904960, η^2^ = 0.902466; post-hoc: Lo vs spontaneous p = 0.60796). However, the mean of peak speed for evoked responses elicited by Hi stimulus strength is significantly larger than that of both Lo stimulus level and spontaneous motifs (Fig. 4D, Hi vs Lo p = 0.0028434, Hi vs spontaneous p = 0.0045889). These results thus suggest that in awake mice, the propagation speed of sensory-evoked activity increases with increasing stimulus levels (similar to anesthetized mice). Also, spontaneous activity propagates at a speed similar to the speed of sensory-evoked activity elicited by Lo stimulus level.

**Fig. 4.**
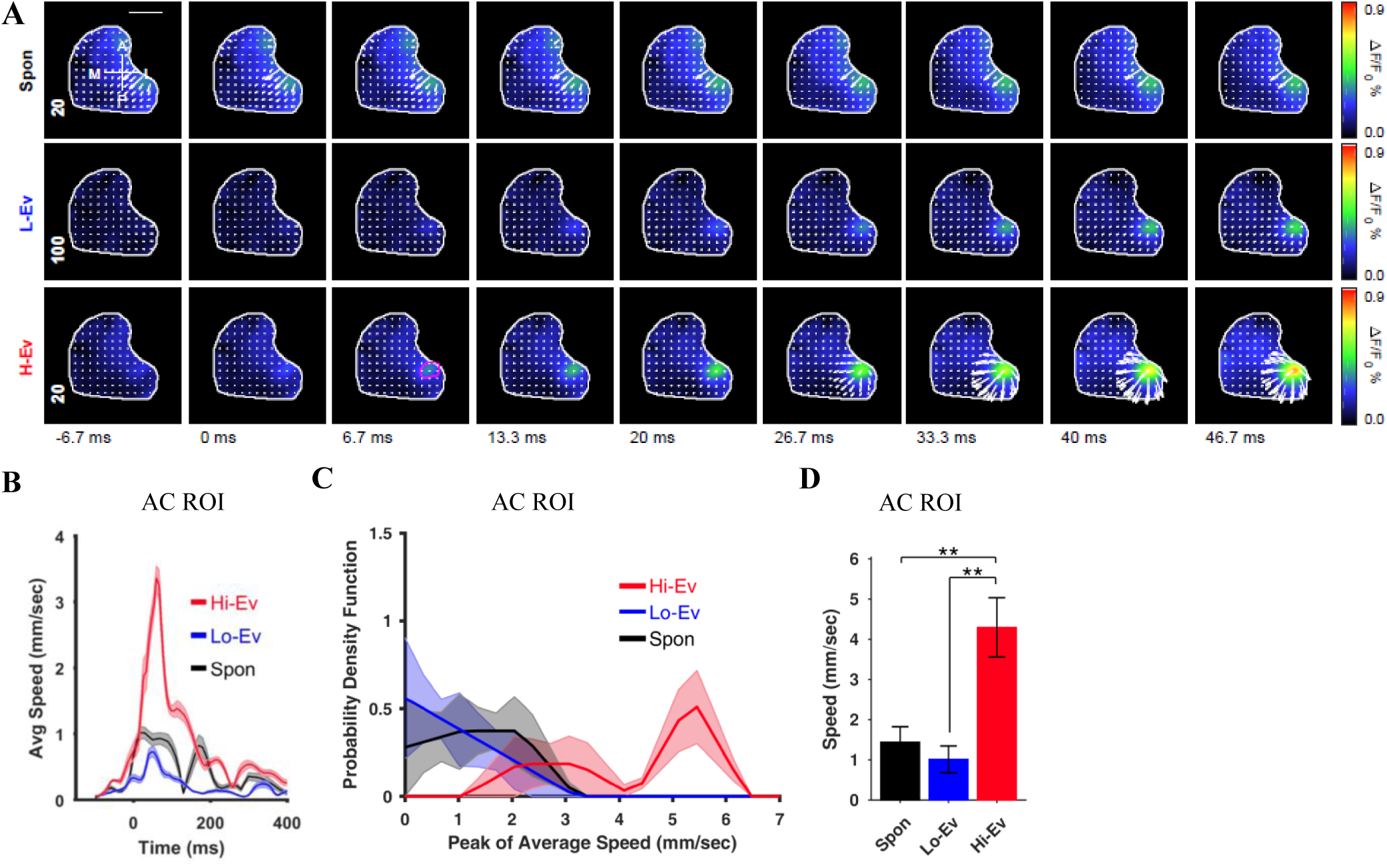
Propagation speed of auditory tone-evoked and spontaneous activity in awake mice. **(A)** Montages of representative examples of spontaneous (top row) and averaged evoked activity in response to auditory stimulation with low (Lo) and high (Hi) stimuli strengths (two bottom rows) with overlaid velocity vector fields determined by the optical flow analysis. Primary AC ROI is outlined in the third column of the bottom row. Compass lines indicate anterior (A), posterior (P), medial (M) and lateral (L) directions. Scale bar is 1 mm. Numbers in the first column indicate scale factor for drawing the velocity vector fields. **(B)** Plots of the average speed signal in the AC-ROI (n=5 animals). The lines represent the mean while shaded regions indicate the SEM across animals. **(C)** Average distributions of peak average speed in the AC-ROI from 5 animals (normalized to the percentage of occurrences). Shaded regions show the SEM across animals. **(D)** Mean±SEM values of peak of average speeds for the AC ROI. ** indicate p<0.01, repeated measure ANOVA with post-hoc Tukey-Kramer correction.

### 3.3 Spatiotemporal propagation patterns of spontaneous activity is more complex than those of sensory-evoked activity in both anesthetized and awake mice

Interaction between neuronal populations is essential for adequate brain function. This interaction often requires brain activity to flow over different brain regions. As demonstrated by wide-field imaging with VSD or glutamate, there is a sequential activation of neuronal populations which have been termed as traveling waves – see review (Muller et al., 2018). We observed propagation patterns of traveling waves and asked whether stimulus levels would affect propagation patterns of sensory-evoked activity and whether these evoked patterns were similar or different from propagation patterns of spontaneous activity. Given that there is a known cortical network responsive to saliency (Downar et al., 2002) and stronger stimulus results in higher saliency (Schultz, 2016), we hypothesized that with increasing stimulus levels, evoked activity would converge to a common propagation pattern. Furthermore, based on the functional roles of spontaneous activity such as memory formation, consolidation, and retrieval, and information processing for carrying out endogenous body functions, we hypothesized that propagation patterns of spontaneous activity would be more complex than those of sensory-evoked activity i.e. spontaneous activity would have a larger repertoire of propagation patterns. To test these hypotheses, we performed two types of analyses. First, we analyzed the distributions of directions at peak values of average speeds of cortical activity in different ROIs using optical flow analysis. The directions were determined from the average velocity vectors. Second, we calculated the trajectory space of cortical activity by building normalized histograms of trajectories. For sensory-evoked and spontaneous activity, the trajectories of responses in all trials and in all motifs were used respectively.

In anesthetized mice, we observed a gradual convergence to a stereotypical direction of propagation with increasing stimulus levels for sensory forelimb evoked responses (Fig. 5). The distributions of directions of velocity vectors of forelimb evoked activity elicited with Lo, Med, and Hi stimulus levels became tighter and the angles became closer to the medio-caudal direction. This result was evident for directions observed in the FLS1 ROI (Fig. 5A) but was more prominent for HLS1 and PTA ROIs (Fig. 5C and 5E respectively). In order to quantify the variability of directions we determined the magnitude of the average of normalized velocity vectors i.e. first we normalized all velocity vectors such that their magnitude was 1, then we averaged all these vectors and calculated the magnitude of the vector sum. A magnitude of the average vector closer to 0 would represent a large variability while a magnitude closer to 1 would represent presence of more dominant direction. The magnitude of the average vector for Lo, Med, and Hi stimulus levels increased in the FLS1 ROI (Fig. 5B) suggesting that with increasing stimulus levels, propagation directions converge to a dominant direction. This was also observed for the HLS1 and PTA ROIs (Fig. 5D and 5F). This matched our visual observation of forelimb evoked VSD activity where for Med and Hi stimulus levels the response activity travels abruptly in the medio-caudal direction. For forelimb motifs in spontaneous activity, although we hypothesized that the activity propagation would be more complex in terms of distributions of directions (i.e. it would have a uniform distribution with all angles represented equally), we observed that the distributions were skewed with the majority of angles representing the caudolateral or lateral direction in the FLS1, HLS1, and PTA ROIs (Fig. 5A, 5C, and 5E). This suggests that spontaneous activity events might have a dominant direction. Furthermore, the magnitude of the average vector for spontaneous events activity shows no significant difference with the magnitude of the average vector for evoked activity elicited with Lo stimulus level in all three ROIs (Fig. 5B, 5D, and 5F). On the other hand, this measurement for spontaneous motifs is significantly smaller than that of Med and Hi stimulus strength (Fig 5D for HLS1, repeated measures ANOVA: F-value = 14.081650, p = 0.0003093654, η^2^ = 0.778781; post-hoc: Med vs spontaneous p = 0.0066370, Hi vs spontaneous p = 0.0091750; 5F for PTA, repeated measures ANOVA: F-value = 6.166851, p = 0.0088490512, η^2^ = 0.606565; post-hoc: Hi vs spontaneous p = 0.0011688). This observation also asserts the high similarity between spontaneous activity and evoked activity at Lo stimulus level. To investigate the structure of trajectories of evoked and spontaneous activity motifs on the cortical mantle, we analyzed normalized histograms of activity trajectories represented as heat maps (Fig. 5G). Bright and dim colors indicate larger and smaller percentage of activity respectively. For forelimb evoked activity, the trajectory space became smaller with increasing stimulus levels (Fig. 5G for representative animal). In contrast, the trajectories for forelimb motifs in spontaneous activity occupied a larger space. For a quantitative comparison of trajectories, we determined the Hausdorff fractal dimension which measures the roughness of the trajectory path. The Hausdorff fractal dimensions of heat maps for spontaneous activity events were significantly larger than those of evoked activity (Fig. 5H, repeated measures ANOVA: F-value = 11.937434, p = 0.0111830599, η^2^ = 0.749019; post-hoc: Med vs spontaneous p = 0.015937). To compare the trajectory space of spontaneous activity events with all levels of evoked responses, the distributions of Hausdorff fractal dimension for both spontaneous motifs and evoked responses pooled from all stimulus intensity levels were generated using a bootstrapping technique (see methods). Fig. 5I shows these distributions for individual animals as well as the mean distributions. The Hausdorff fractal dimension distribution of spontaneous motifs spans to higher values and has higher mean value compared to the one from evoked responses (Wilcoxon rank sum test on the mean values for animals, p = 0.047619, n = 5). A larger Hausdorff fractal dimension corresponds to a larger space occupancy for trajectories (Singh et al., 2016) and thus spontaneous activity has a larger spatial presence as compared to sensory-evoked activity. For hindlimb stimulation, the distributions of directions observed in the HLS1 ROI for Lo, Med, and Hi, stimulus levels were similar with comparable average vector magnitudes (Fig. S5A and S5B). In the FLS1 and PTA ROIs however, the distributions of directions converged with increasing stimulus levels suggesting a stereotypical expansion of hindlimb evoked activity with Hi stimulus strength (Fig. S5C and S5E). The hindlimb motifs in spontaneous activity, similar to forelimb, had the majority of propagation angles in the caudolateral and lateral directions in the HLS1, FLS1, and PTA ROIs. The average vector magnitudes tend to be smaller than evoked responses at all levels (Fig. S5). The Hausdorff fractal dimensions of trajectory heat maps for hindlimb motifs in spontaneous activity were significantly larger than those of hindlimb-evoked activity (Fig. S5G and S5H, repeated measures ANOVA: F-value = 8.356785, p = 0.0028671496, η^2^ = 0.676291; post-hoc: spontaneous vs Med p = 0.00083979, : spontaneous vs Hi p = 0.014265). For the visual sensory modality, we observed a stereotypical distribution of directions for evoked activity towards the anterior brain regions while spontaneous activity events traveled in all directions (Fig. S6A and S6B). Similar to what was observed for the forelimb and hindlimb somatosensory modalities, the Hausdorff fractal dimension was larger for the spontaneous activity heat maps (Fig. S6C and S6D, paired sample t-test p = 0.0087369698). These results combined suggest that spontaneous activity has more complex spatiotemporal propagation patterns with a wider range of angles and larger trajectory spaces than evoked activity elicited with Hi stimulus levels. Although, the distributions of directions of velocity vectors suggest that spontaneous activity motifs might have a dominant direction, its average vector magnitude is similar to Lo stimulus level and is significantly smaller than for higher stimulus strengths. Alternatively, evoked activity converges to a common propagation pattern with increasing stimulus levels. Finally, spontaneous motifs occupy a larger trajectory space compared to evoked responses.

**Fig. 5.**
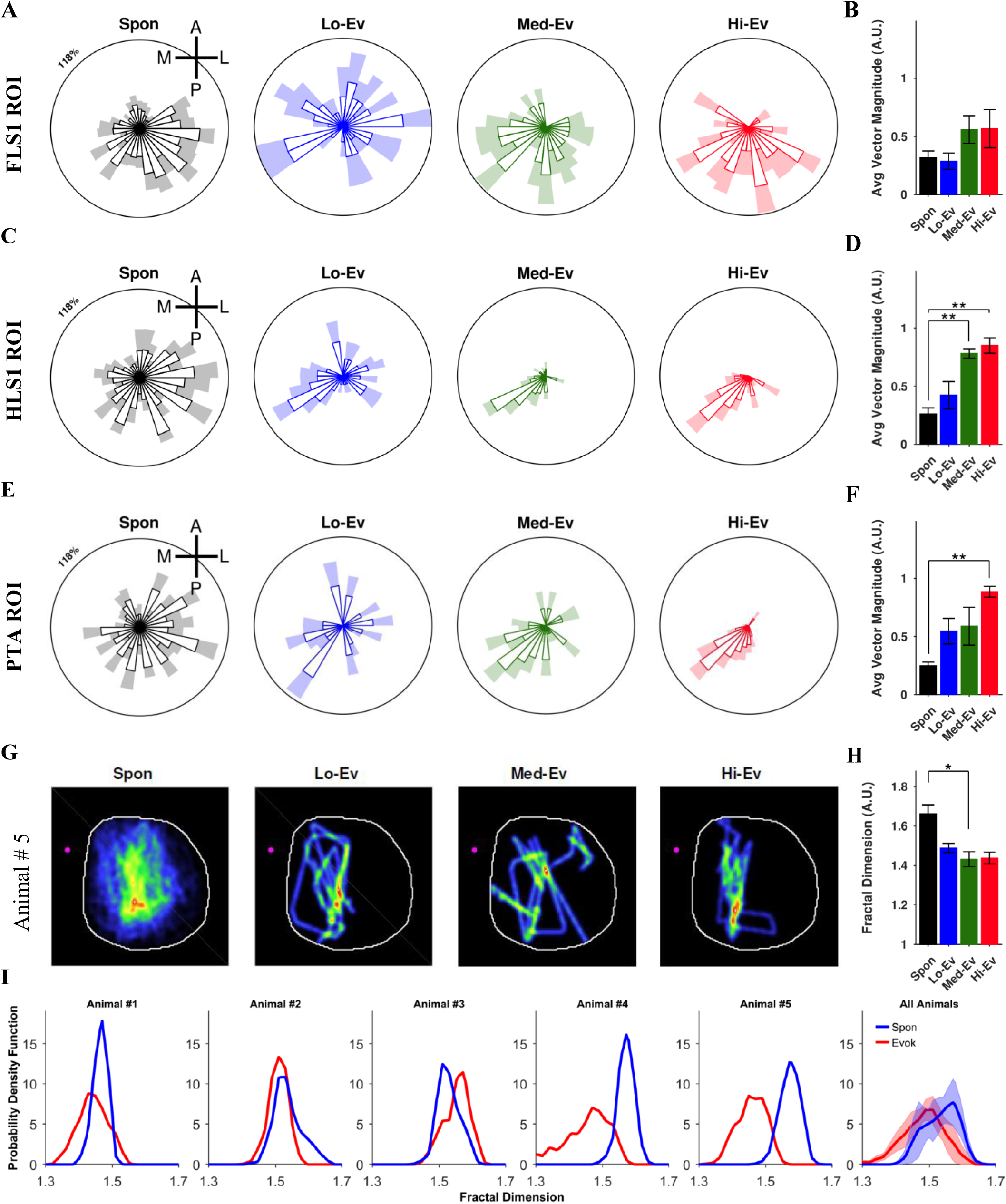
Propagation patterns of forelimb-evoked activity converge as stimulus levels increases and are less complex than those of spontaneous activity in anesthetized mice. **(A)** Average distributions of directions of the peak velocity vectors in the FLS1 ROI for spontaneous (Spon) activity and evoked (Ev) activity elicited with Lo, Med, and Hi stimulus levels normalized to the percentage of occurrences across animals (n=5). **(B)** Magnitude of the average of normalized velocity vectors. (**C**-**D)** and (**E**-**F)** similar to (**A**-**B**) but for HLS1 and PTA ROIs respectively. **(G)** Normalized histogram of activity trajectories represented as a heat map with warm and cold colors indicating larger and smaller occurrences of activity passing through a given point on the cortical surface. **(H)** Mean±SEM values of Hausdorff fractal dimension of heat maps. **(I)** Distributions of the Hausdorff fractal dimension for evoked and spontaneous activity for each animal and average for all animals (last column). * and ** indicate p<0.05 and p<0.01 respectively, repeated measure ANOVA with post-hoc Tukey-Kramer correction.

Next, we investigated the propagation patterns of spontaneous and evoked activity in awake mice. For this analysis, we selected two regions over the mouse cortical surface based on visual observation of trajectories of auditory stimulation responses. These regions are adjacent to auditory cortical areas, namely anteromedial to AC ROI (AC-AM) near barrel cortex and posteromedial to AC ROI (AC-PM) near visual cortex. In the selected ROIs, results were similar to the ones observed for anesthetized mice. Sensory-evoked activity converged to a propagation pattern with increasing stimulus levels as suggested by tendency to increase in magnitude of the average of normalized velocity vectors for Lo and Hi stimulus levels (Fig. 6B, 6D, and 6F). However, the magnitudes are less than 0.5 (i.e. closer to 0), suggesting a large variability in the distributions of directions, which is evident in Fig. 6A. This matches our visual observation of auditory-evoked activity originating and expanding in the AC ROI thus giving rise to velocity vectors in all directions. In the AC-AM ROI (Fig. 6C and 6D) however, the activity propagation patterns were not as clear either because the evoked activity does not travel strongly to this ROI or because when the animals are awake, the evoked activity co-exists with spontaneous activity and becomes less salient compared to the anesthetized case. For AC-PM ROI, it is interesting to note that evoked activity with Hi stimulus level has a tight distribution of directions (Fig. 6E). This suggests that with Hi stimulus strength, evoked activity traveled from the AC to AC-PM ROI in a stereotypical fashion across animals. For spontaneous activity motifs in auditory region, we observed dominant directions towards the midline of the cortical surface (Fig. 6A, 6C, and 6E). The magnitude of average velocity vector is less than 0.5, which also suggests a large variability in the direction of spontaneous activity events (Fig. 6B, 6D, and 6F). In AC-PM ROI there is no significant difference between the velocity vectors average for Lo and Hi stimulus levels and spontaneous motifs, but Hi stimulus level tends to show more stereotyped propagation direction. As described above, here also, we calculated heat maps to study the structure of cortical activity (Fig. 6G). As observed for anesthetized mice, the trajectory space occupied by spontaneous activity events was larger than that occupied by Hi stimulus evoked activity, which is confirmed by a significantly larger Hausdorff fractal dimension for spontaneous activity (Fig. 6H, repeated measures ANOVA: F-value = 11.565240, p = 0.0043613067, η^2^ = 0.743017; post-hoc: spontaneous vs Hi p = 0.0028885). Finally, comparing distributions of Hausdorff fractal dimension for spontaneous motifs and awake evoked responses pooled from both Lo and Hi stimulus levels indicate that spontaneous motifs exhibit larger values (Wilcoxon rank sum test on the mean values for animals, p = 0.047619, n = 5).

**Fig. 6.**
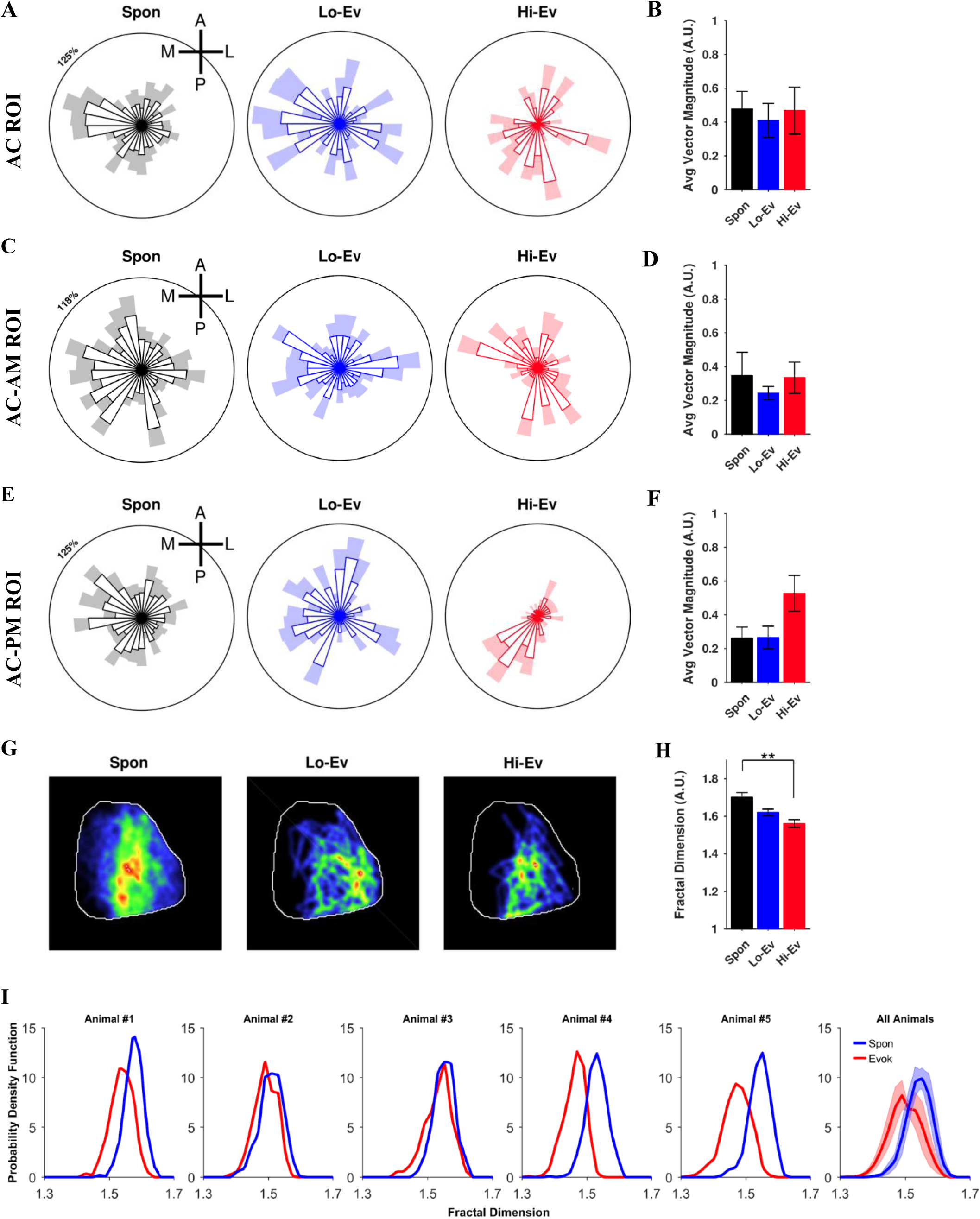
Propagation patterns of auditory tone-evoked activity converge with increasing stimulus levels and are less complex than those of spontaneous activity in awake mice. **(A)** Average distributions of the directions of the peak velocity vectors in the AC ROI for spontaneous (Spon) activity and evoked (Ev) activity elicited with Lo and Hi stimulus levels normalized to the percentage of occurrences over animals (n=5). **(B)** Magnitude of the average of normalized velocity vectors. **(C** and **D)** and (**E** and **F**) similar to (**A** and **B**) but for AC-AM and AC-PM ROIs respectively. **(G)** Normalized histogram of activity trajectories represented as a heat map with warm and cold colors indicating larger and smaller occurrences of activity passing through a given point on the cortical surface. **(H)** Bar plot of the mean±SEM of the Hausdorff fractal dimension of the heat maps. **(I)** Distributions of the Hausdorff fractal dimension for evoked and spontaneous activity for each animal and average for all animals (last column). ** indicates p<0.01, repeated measure ANOVA with post-hoc Tukey-Kramer correction.

**Fig. 7.**
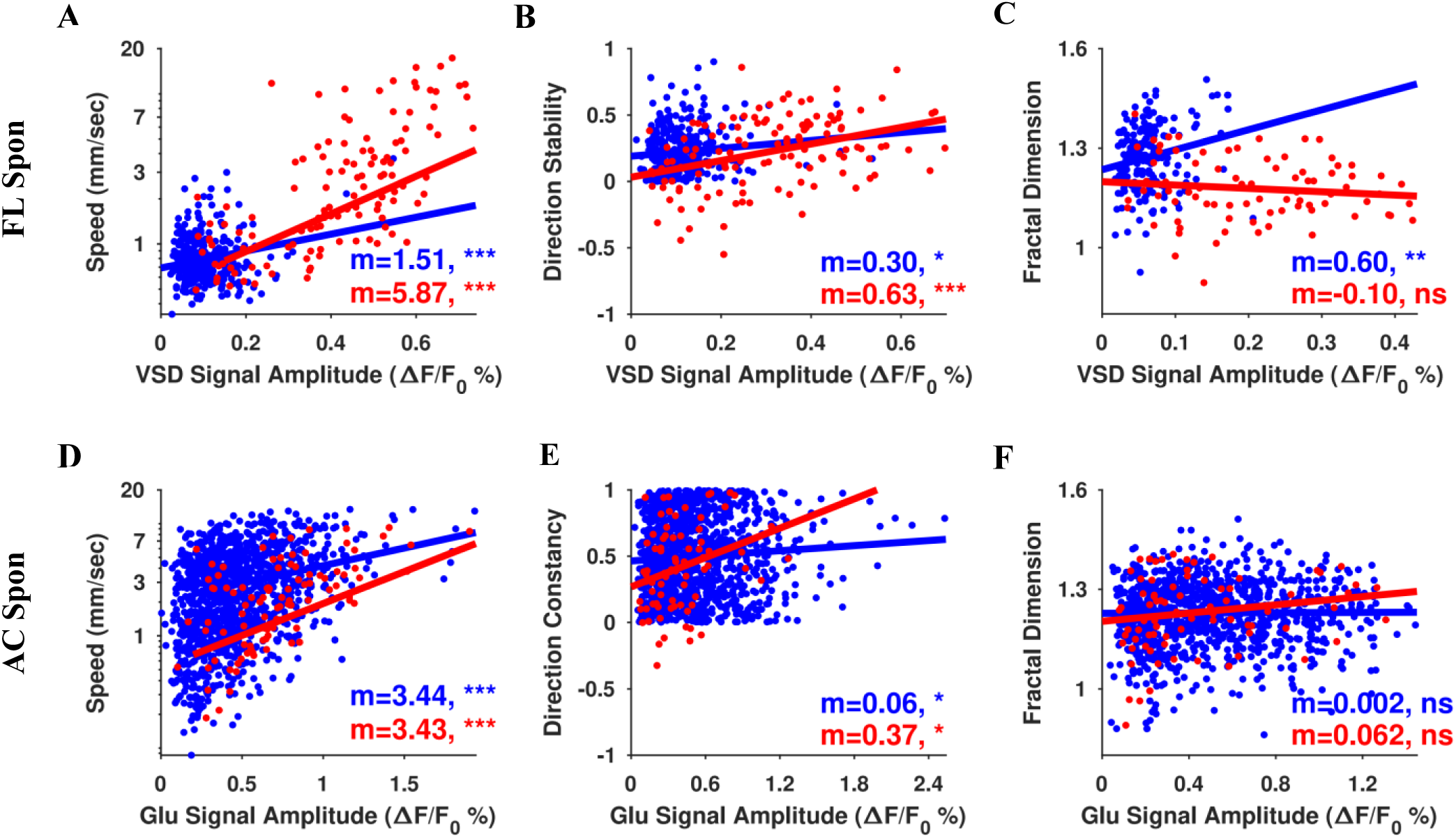
Speed, flow direction stability, and fractal dimension of the propagation of spontaneous and evoked activity are positively correlated with its amplitude in awake and anaesthetized mice. **(A)** Scatter plot of VSD signal amplitudes for forelimb spontaneous motifs (blue) and evoked (red) activity versus peak velocity vectors in the FLS1 ROI for anaesthetized mice. Each dot represents one trial. The data is pooled from all animals (n=5). The lines represent linear regression models fitted to the corresponding data. **(B)** Scatter plot and linear regression fit for VSD signal amplitude for forelimb spontaneous (blue) motifs and evoked activity (red) in the HLS1 ROI vs flow direction stability for anesthetized mice. **(C)** Similar to **(A)** but for VSD signal amplitudes in the entire imaging window vs fractal dimensions of spontaneous motifs (blue) and evoked activity (red). (**D-F**) Similar to (**A-C**) but for auditory spontaneous motifs (blue) and evoked activity in AC in awake mice. The data is pooled from all animals (n=5). m is the estimated slope. *, **, and *** indicates p<0.05, p<0.01, and p<0.001 respectively, for the p-value of the linear regression slope. Note that speed axis in **A** and **D** is logarithmic scale.

Altogether, our results suggest that the propagation patterns of spontaneous activity are more complex by occupying larger trajectory spaces compared to those of evoked activity in awake mice.

### 3.4 Propagation speed of spontaneous activity is positively correlated with its amplitude while its pattern of propagation and direction stabilize as amplitude increases

As reported earlier, we found that the amplitude of evoked activity is positively correlated with its speed and also with the stability of its propagation. Here, we asked if the propagation speed, direction, and pattern of spontaneous activity was also correlated with its amplitude. We found that for evoked and spontaneous activity in the forelimb and auditory regions, the propagation speed was positively correlated with its amplitude (scatter plots in Fig. 7A and D where each blue dot represents a single spontaneous activity motif and each red dot represent a single evoked trial). The fitted linear regression model showed a positive slope (Fig. 7A, spontaneous: linear coefficient=1.509 and p-value=0.000057533; evoked: linear coefficient=5.869 and p-value=3.5064×10^−11^, and Fig. 7D, spontaneous: linear coefficient=3.437 and p-value=1.2669×10^−38^; evoked: linear coefficient=3.431 and p-value=9.8767×10^−10^). A similar relationship was also found between propagation direction stability and amplitude (Fig. 7B, spontaneous: linear coefficient=0.296 and p-value=0.047929; evoked: linear coefficient=0.626 and p-value=0.000019484, and Fig. 7E, spontaneous: linear coefficient=3.437 and p-value=1.2669×10^−38^; evoked: linear coefficient=3.431 and p-value=9.8767×10^−10^). The stability of propagation pattern measured as the Hausdorff fractal dimension however was more stable with increasing amplitude for spontaneous activity in the forelimb region compared to auditory region (Fig. 7C, spontaneous: linear coefficient=0.601 and p-value=0.0046286; evoked: linear coefficient=-0.101 and p-value=0.31972, and Fig. 7F, spontaneous: linear coefficient=0.002 and p-value=0.83653; evoked: linear coefficient=0.062 and p-value=0.13361).

Similarly spontaneous activity in the forelimb region show significant positive correlation with amplitude and speed (Fig. S7A, spontaneous: linear coefficient=0.539 and p-value=0.021671; evoked: linear coefficient=5.725 and p-value= 0.000027819) stability of propagation direction in the PTA ROI (Fig. S7B, spontaneous: linear coefficient=0.336 and p-value=0.0066435; evoked: linear coefficient=0.412 and p-value=0.025690) and fractal dimension of patterns in the entire imaging field (Fig. S7C, spontaneous: linear coefficient=0.622 and p-value=0.0000023820; evoked: linear coefficient=0.057 and p-value=0.77803). For spontaneous motifs in the visual region and the whole imaging cortical area all three measurements were positively correlated with the amplitude (Fig. S7D, spontaneous: linear coefficient=6.038 and p-value=2.6386×10^−38^; evoked: linear coefficient=0.057 and p-value=1.4251×10^−60^. Fig. S7E, spontaneous: linear coefficient=0.389 and p-value= 2.4717×10^−8^; evoked: linear coefficient=0.210 and p-value= 0.0018673. Fig. S7F, spontaneous: linear coefficient=0.202 and p-value=0.0014744; evoked: linear coefficient=0.110 and p-value=0.017036).

## 4 Discussion

In this paper, we used voltage-sensitive dye and glutamate imaging in anesthetized and awake mice respectively to characterize cortical dynamics of spontaneous and evoked activity at the mesoscale level. Our main conclusions were similar in anesthetized and awake mice preparations, i.e., sensory-evoked activity and spontaneous activity are similar to each other for low stimulus strength but as the stimulus strength increases, they become different from each other. Previous studies have compared spontaneous and evoked activity using different parameters such as the temporal structure of single-unit activity, unit activity correlations, or dimensionality. In contrast, in this study we use more direct parameters of cortical dynamics such as amplitude, direction and speed of propagation, and complexity of the propagation trajectories. This more direct characterization of the cortical dynamics might be less influenced by confounding effects introduced in the methods to calculate such more abstract measurements.

### 4.1 The effect of stimulus strength on sensory-evoked activity

When the stimulus strength was increased, the amplitude, spread, and speed of the evoked cortical response increased. These results are consistent with previous studies conducted to understand the relationship between cortical responsiveness, and the strength of sensory stimulation (Jancke et al., 1998; Spenger et al., 2000; Peeters et al., 2001; Petersen et al., 2003a; Muniak et al., 2007). The positive correlation between stimulus strength and amplitude, spread and speed of evoked cortical activity could be explained in two ways. First, since a larger number of subcortical neurons are activated with stronger stimulus, these networks, in turn, activate a larger region of cortex through one-to-one connections (Jones, 1998; Sato et al., 2012). Alternatively, the larger spatial spread of evoked cortical response in the cortex might be attributed to enhanced activation of cortico-cortical networks driven by stronger localized subcortical inputs. Further experiments are required to investigate the subcortical-cortical circuit loops involved in shaping the extent of cortical activation (Jones, 1998). In addition to the relationship between the extent of activity propagation and stimulus intensity, our results also suggest a positive correlation between the speed of the activity propagation and the stimulus strength. However, we found a negative correlation between the stimulus strength and the variability in the direction of activity propagation. Similar observations have been reported in psychophysics and electrophysiological research, where the relationship between stimulus strength or contrast and speed, accuracy or variability of response are negatively correlated (Palmer et al., 2005; Deco and Hugues, 2012; Sadagopan and Ferster, 2012). An alternative explanation for observing a larger variability of directions for weaker stimuli compared to when giving stronger stimulation might be attributed to a less accurate detection of propagation direction by the optical flow method used in this study.

### 4.2 Comparison between sensory-evoked and spontaneous activity patterns

Our results indicate that the amplitude of sensory-evoked cortical response is larger than the amplitude of spontaneous activity events for larger stimulus intensity (Med and Hi) in the somatosensory and visual cortices of anesthetized mice and auditory cortex of awake mice. A related comparison has been previously reported for the visual cortex of anesthetized cats and monkeys using electrophysiological recordings (Nauhaus et al., 2009). In their work, low contrast stimulus yielded high amplitude LFP waves that travelled longer distances than high-contrast evoked waves. In addition, it has been reported that in V1 of anesthetized cats, the amplitude of VSD signals of evoked and spontaneous activity are comparable (Grinvald et al., 2003). The discrepancy between these studies and our results might be due to the level of stimulus strength used, or scale (e.g., field of view) at which cortical activity was measured (micro for electrophysiology vs meso for wide-field VSD), or the origin of the signal (mostly subthreshold membrane depolarization for VSD vs a combination of subthreshold and suprathreshold activity for LFP signal) (Grinvald and Hildesheim, 2004).

We also found that on average, sensory-evoked activity travels faster than the spontaneous activity. On average, evoked activity in the FLS1 area travels at 3.35±0.63 mm/s while forelimb motifs in spontaneous activity travels at 0.91±0.08 mm/s. For hindlimb stimulation and HLS1 area, evoked activity travels at 2.37±0.43 mm/s and hindlimb spontaneous motifs travel at 0.88±0.08 mm/s. Sensory evoked activity in the visual cortex and its spontaneous motifs travels at 4.79±1.88 mm/s and 1.92±0.55 mm/s respectively. It is important to mention that in previous VSD recordings in the visual cortex of anesthetized rats, it is reported that spontaneous activity travels faster than evoked activity (16 mm/s and 10 mm/s respectively) (Han et al., 2008). In contrast, we found that with our method, this is not the case for the somatosensory cortex, nor for visual cortex. Moreover, (Nauhaus et al., 2009; Sato et al., 2012) reported that in anesthetized monkeys and cats, spontaneous activity travels at ∼0.3m/s in visual cortex using electrophysiological studies. One reason for the discrepancy between the speeds reported in these studies and our results might be the differences in methodology used to quantify the propagation of activity in VSD recordings and the scale of the neuronal population and brain size from which the activity was measured. In terms of propagation, sensory-evoked cortical activity has more stereotyped directions of propagation and trajectories (with higher stimulus strengths) compared to spontaneous activity which shows a broader distribution of possible directions of activity propagation. We also show that the trajectories of spontaneous activity sample a broader space than during evoked activity.

### 4.3 Possible mechanisms for the differences between sensory-evoked and spontaneous activity

Why would evoked activity have larger amplitudes than spontaneous activity? And why would spontaneous activity travel slower and longer distances than evoked activity? The explanation for this might lie in the difference in their origin and the particular circuitry of the laminar structure of the cortex. While spontaneous activity spreads upward from deep layers and slowly across columns, evoked activity, on the other hand, initiates in thalamocortical recipient layers and spreads rapidly across columns (Sakata and Harris, 2009; Chauvette et al., 2010; Beltramo et al., 2013; Petersen and Crochet, 2013).

Since in VSD, most of the signal comes from superficial cortical layers that contain dendritic trees of neurons located in both layer 2/3 (L2/3) and layer 5 (L5) (Chemla and Chavane, 2010; Mohajerani et al., 2011), the explanation for the differences reported in this work might be found by revisiting the input circuitry of L2/3 of the cortex (Sakata and Harris, 2009). According to the canonical circuit, sensory evoked activity flows into and through the cortex first from the thalamus to layer 4 (L4), then to L2/3 and finally to L5/6 (Douglas and Martin, 2004; DeNardo et al., 2015). Although it has been shown that the connectivity of the somatosensory cortex largely follows this canonical circuit, there is recent evidence that L2/3, L5, and L6 also receive inputs directly from the thalamus (Viaene et al., 2011; Constantinople and Bruno, 2013; DeNardo et al., 2015). Thus, the larger amplitude and faster propagation observed during evoked activity could be due to strong thalamic inputs apart from the already existent cortico-cortical connections used during spontaneous activity (Bruno and Sakmann, 2006). Finally, a factor for the shorter propagation of evoked activity compared to spontaneous activity might be due to the inhibition triggered by the thalamocortical system engaged after the initiation of evoked activity (Sheroziya and Timofeev, 2014; McCormick et al., 2015).

The observation that spontaneous activity has a larger number of patterns compared to evoked activity in a sensory region might be explained by how networks are wired during early and later development. It is known that during early development, neuronal connections are first formed by genetically driven molecular factors and later shaped by patterned spontaneous activity so much so that networks can interpret sensory signals or show tuning without prior experience e.g., presence of orientation selectivity in the visual system before visual experience (Chapman et al., 1999; Ackman and Crair, 2014; McVea et al., 2016), path integration during early stages before maturation of grid cells (Bjerknes et al., 2018). In a sensory brain region therefore, during early development one would expect to see a larger repertoire of spontaneous activity patterns compared to evoked activity which perhaps is carried forward to adulthood (McVea et al., 2017). Furthermore, in an adult sensory brain region e.g., visual cortex, one might expect to see neuronal activity correlated with many non-visual behaviors such as those associated with facial motion (Stringer et al., 2019). Therefore, during development, primary sensory areas also get wired with many other brain regions. In studies such as ours where behavior is not recorded and quantified, activity associated with non-sensory behaviors would be deemed spontaneous activity (at least in awake preparation) and therefore would have more diverse patterns than sensory-evoked activity.

### 4.4 Functional relevance of differences between sensory-evoked and spontaneous activity

While evoked activity is arguably only involved in sensory processing, spontaneous activity has multiple functions including memory consolidation, refinement of cortical circuitry, and synaptic homeostasis. Therefore, the possible realm of computations involved during the communication amongst different cortical circuits during spontaneous activity might be broader than the ones involved only in processing sensory information (Luczak et al., 2007). Moreover, the differences in speed and propagation can be strongly related to the functional roles of spontaneous and evoked activity in learning and memory (Mercado, 2014). These observations are based on evidence about the sparse action potential firing of L2/3 and which, in combination with its dense subthreshold inputs can be exploited for associative learning (Petersen and Crochet, 2013). This sparse action potential firing of L2/3 might be paired with specific sensory events, top-down input, and neuromodulatory input to regulate plasticity. Therefore, the temporal differences between spontaneous and evoked activity, in the order of milliseconds, might be directly related to synaptic processes. For example, the faster propagation of evoked activity might involve different synaptic processes to facilitate the encoding of relevant events. On the other hand, the longer propagation of spontaneous activity might involve the ‘replay’ of coherent experiences or memories distributed over large cortical areas for consolidation (Wilson and McNaughton, 1994; Hoffman et al., 2007; Ji and Wilson, 2007). In contrast, the similarities found between spontaneous activity and sensory-evoked activity with low stimulus strength at both micro and meso scales, might be due to the restrictions imposed by the brain circuitry (Luczak and Maclean, 2012) i.e., the underlying common brain hardware.

This study clarifies the discrepancies in similarities and differences previously reported for spontaneous and evoked activity. Using wide-field imaging allowed for better resolution in space than previous electrophysiological studies (Luczak et al., 2009; Sakata and Harris, 2009) and in time than previous fMRI studies (Jancke et al., 1998; Spenger et al., 2000; Huang et al., 2015), or imaging studies over a small cortical area (Han et al., 2008). Spontaneous activity characteristics including amplitude, propagation speed, and direction distribution is similar to evoked activity elicited with lower stimulus strength. Also, the trajectory space of spontaneous activity covers a larger cortical area compared to evoked activity.

### 4.5 Differences in spontaneous and evoked activity as a biomarker for neuropsychiatric disorders

Alterations in the balance of excitation and inhibition ratio in the cerebral cortex have been suggested as an explanation for various neurological and psychiatric disorders such as schizophrenia (Goel and Portera-Cailliau, 2019). Similarly, the differences in sensory-evoked and spontaneous activity might be good candidates as objective biomarkers for clinical diagnosis of neurological and psychiatric disorders. Resting-state spontaneous activity alone measured with fMRI (Fox and Raichle, 2007; Kiviniemi, 2008) has been used to differentiate between normal individuals and patients with various neurological diseases (Fox and Greicius, 2010) such as Alzheimer disease (Zeng et al., 2019), Parkinson’s disease (Zhang et al., 2019), schizophrenia (Nejad et al., 2012), tinnitus (Cai et al., 2019), unilateral amblyopia (Dai et al., 2019), amyotrophic lateral sclerosis (Luo et al., 2012), idiopathic trigeminal neuralgia (Yuan et al., 2018), and corneal ulcer (Shi et al., 2019). In addition, combined task-evoked and spontaneous activity fMRI have been used for pre-operative mapping (Fox et al., 2016). In the above-mentioned studies, three main types of analysis have been used; 1) regional coherence in which similarity of activity of neighboring voxels is assessed using cross-correlation, 2) power spectrum analysis in which amplitudes of low-frequency fluctuations are measured, and 3) spatial pattern analysis in which the topology of activity is compared between controls and diseased individuals (Fox and Greicius, 2010). These analyses lack focus on the dynamics of activity i.e., speed and propagation. A greater understanding of the parallel time-varying properties of resting-state activity is increasingly believed to be essential to understanding brain function in health and disease). We propose the use of task-independent graded stimuli for recording local evoked activity and its comparison with corresponding local resting-state spontaneous activity for identifying diseased individuals. This may have particular utility in revealing the neural circuit dysfunction underlying altered sensory processing in neuropsychiatric disease contexts. For example, individuals with schizophrenia exhibit deficits in auditory sensory gating as revealed by the suppression of event related potential markers P50 and N100 (Brockhaus-Dumke et al., 2008; Javitt, 2009). The methods described in our study allow the characterization of the dynamics of activity, and therefore, would provide superior discrimination measurements. Moreover, this approach could be extended to human studies using fMRI by adapting our methods to 3D version of the optical flow analysis for estimation of activity speed and propagation (von Tiedemann et al., 2010; Rajna et al., 2019).

### Disclosures

The authors have no relevant financial interests in the manuscript and no other potential conflicts of interest to disclose.

## Acknowledgments

This work was supported by a Natural Sciences and Engineering Research Council of Canada (NSERC) Discovery Grant #40352, Campus Alberta for Innovation Program Chair, Alberta Alzheimer Research Program to MHM and NSERC CREATE in BIP doctoral fellowship to NA. We thank Jianjun Sun for assistance with surgeries, Behroo Mirza Agha and Di Shao for animal husbandry, and Allen Chan and Timothy H. Murphy for critical reading of the manuscript.

## Author Contribution

Conceptualization, M.H.M., N.A., S.I., and E.B.C., A.L., B.L.M.; Methodology, M.H.M., N.A., S.I., and E.B.C.; Formal Analysis, N.A. and S.I.; Writing - Original Draft, M.H.M., N.A., S.I., and E.B.C.; Writing - Review & Editing, M.H.M., N.A., S.I., A.L., E.B.C., and B.L.M.; Funding Acquisition, M.H.M., B.L.M., A.L.; Resources, M.H.M.; Supervision, M.H.M.

