## Supplemental Figures for "Spatiotemporal structure of sensory-evoked and spontaneous activity revealed by mesoscale imaging in anesthetized and awake mice"

**C**

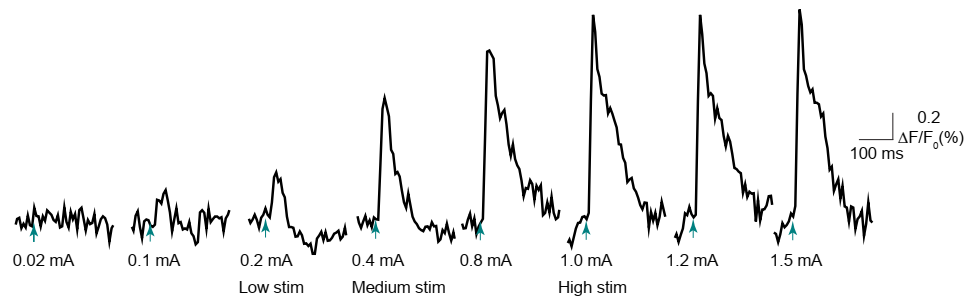

**A**

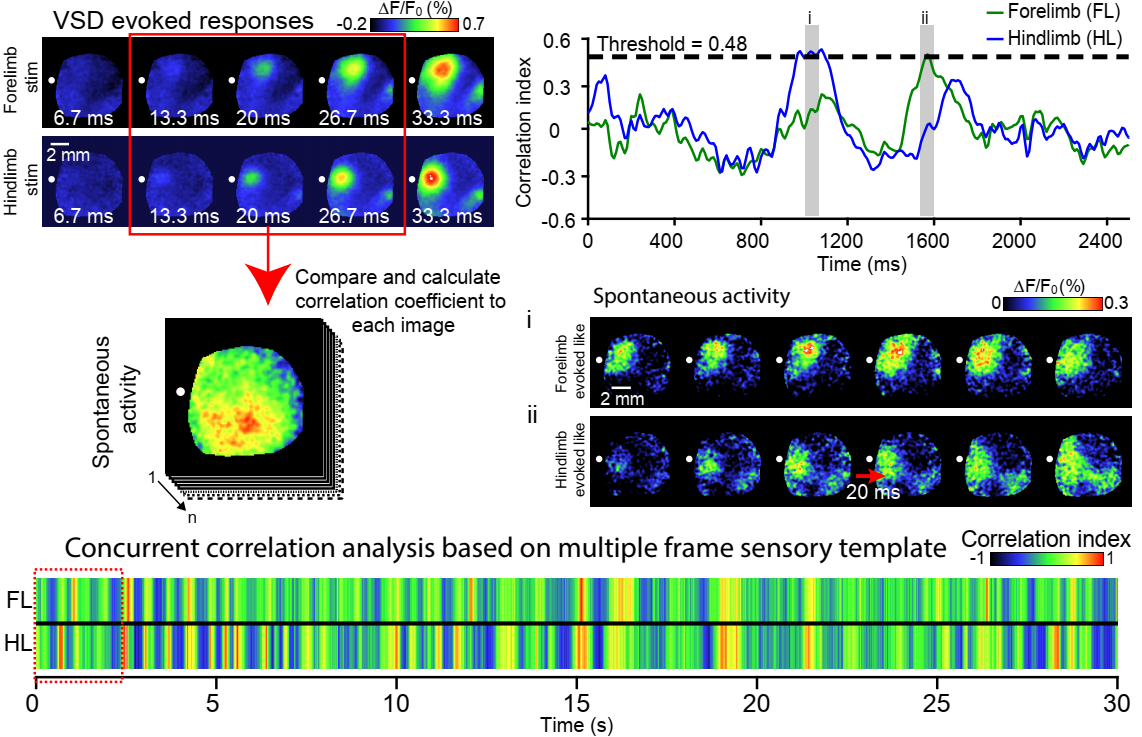

**B**

**D**

**Fig. SM1 Sensory evoked response categorization and spontaneous motifs determination.** **(A)** Forelimb (FL) VSD responses to multiple stimulus strength in a representative animal in the FL ROI. The green arrows show the stimulus onset. Out of all stimulus levels, three levels are selected as low, medium, and high responses. **(B)** (top) Montage of VSD imaging responses to forelimb and hindlimb (HL) stimulation. The red square shows the selected template frames. (bottom) For each modality the template frames were correlated with spontaneous activity frames to determine similar events in spontaneous activity. **(C)** (top) Correlation index values for FL and HL template matching in a 2.5s. The instances that the correlation index passes the threshold are selected as spontaneous motifs. (bottom) FL and HL examples of spontaneous motifs determined. **(D)** FL and HL template matching correlation index for a 30s-time window.

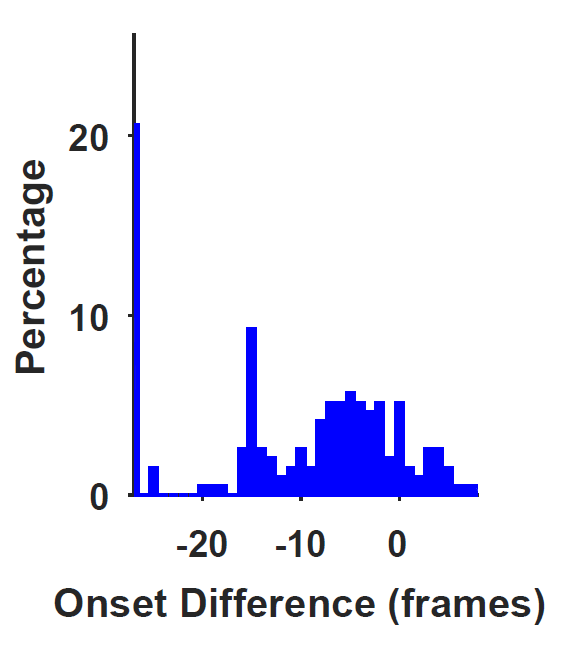

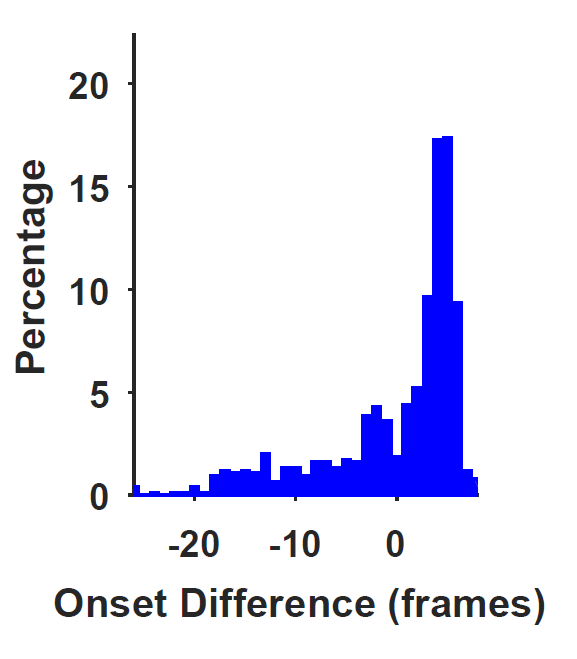

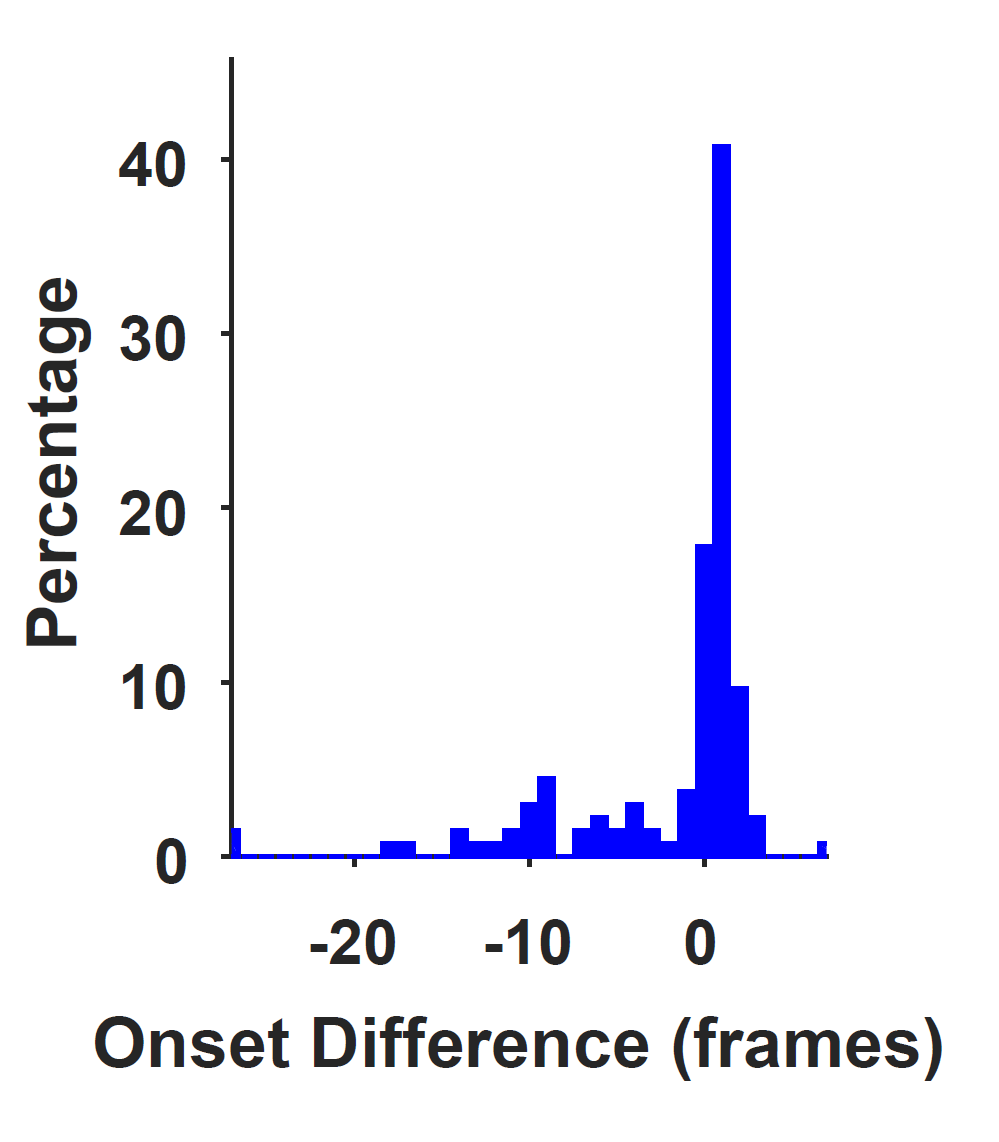

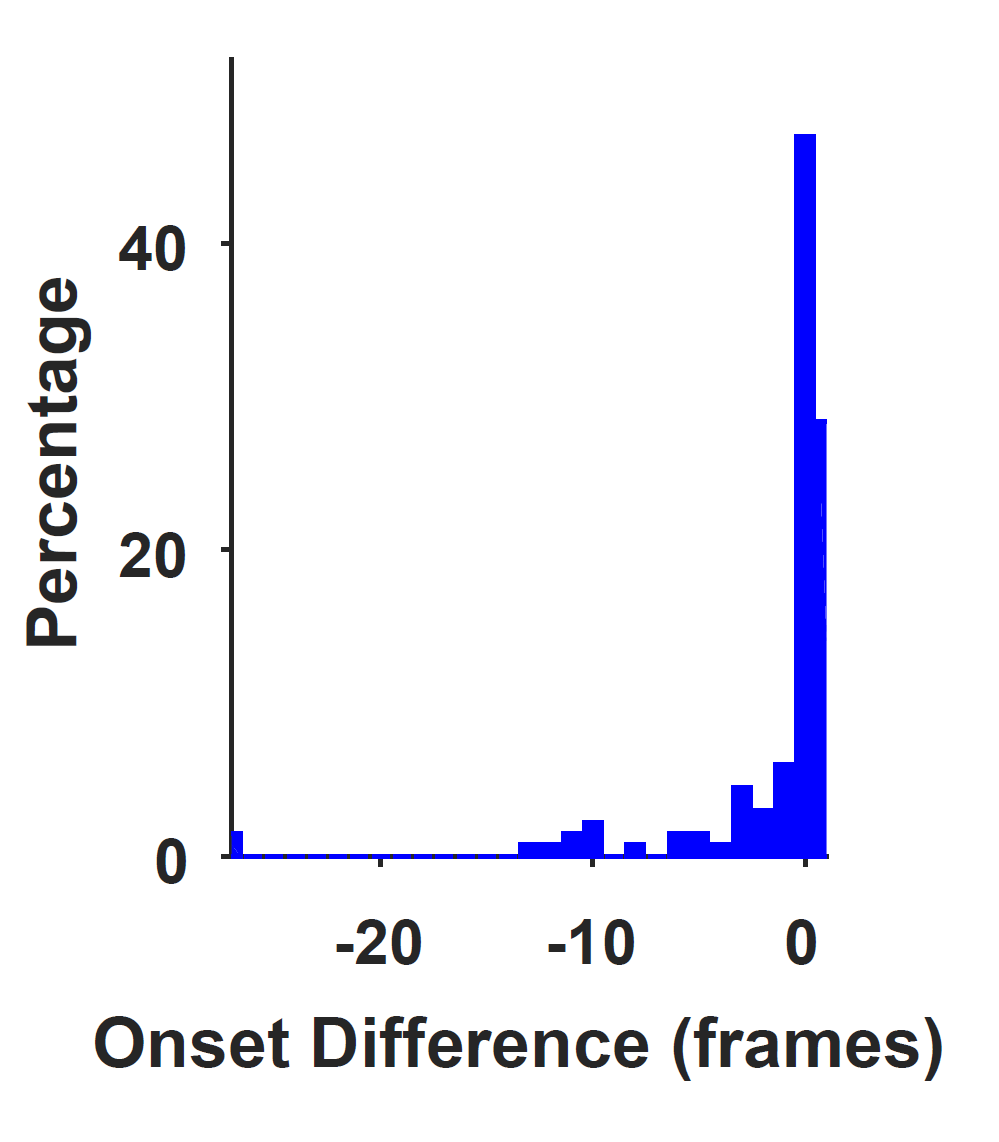

**A**

**B**

**C**

**D**

**E**

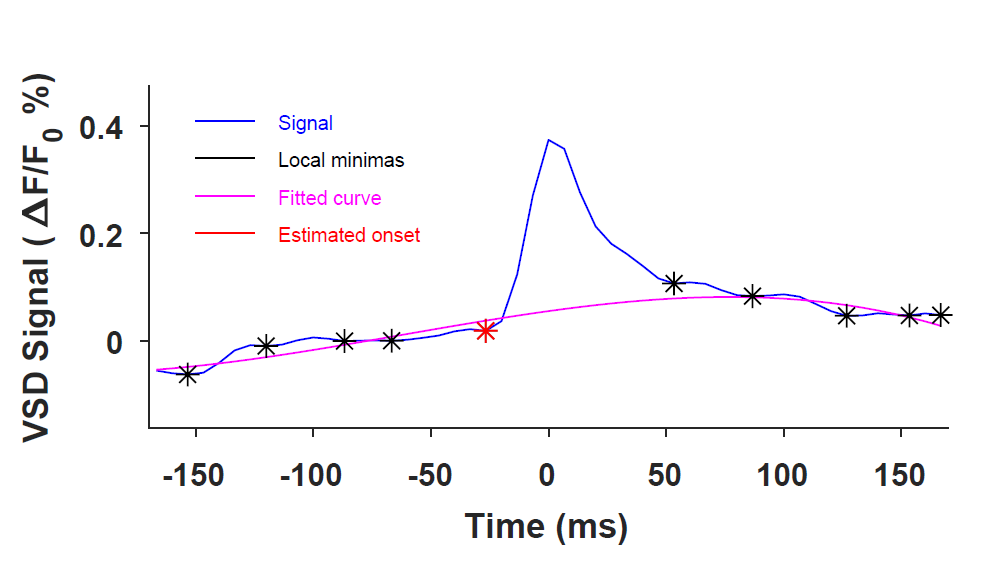

Forelimb evoked

Hindlimb evoked

Visual
evoked

Auditory evoked
(Glutamate Signal)

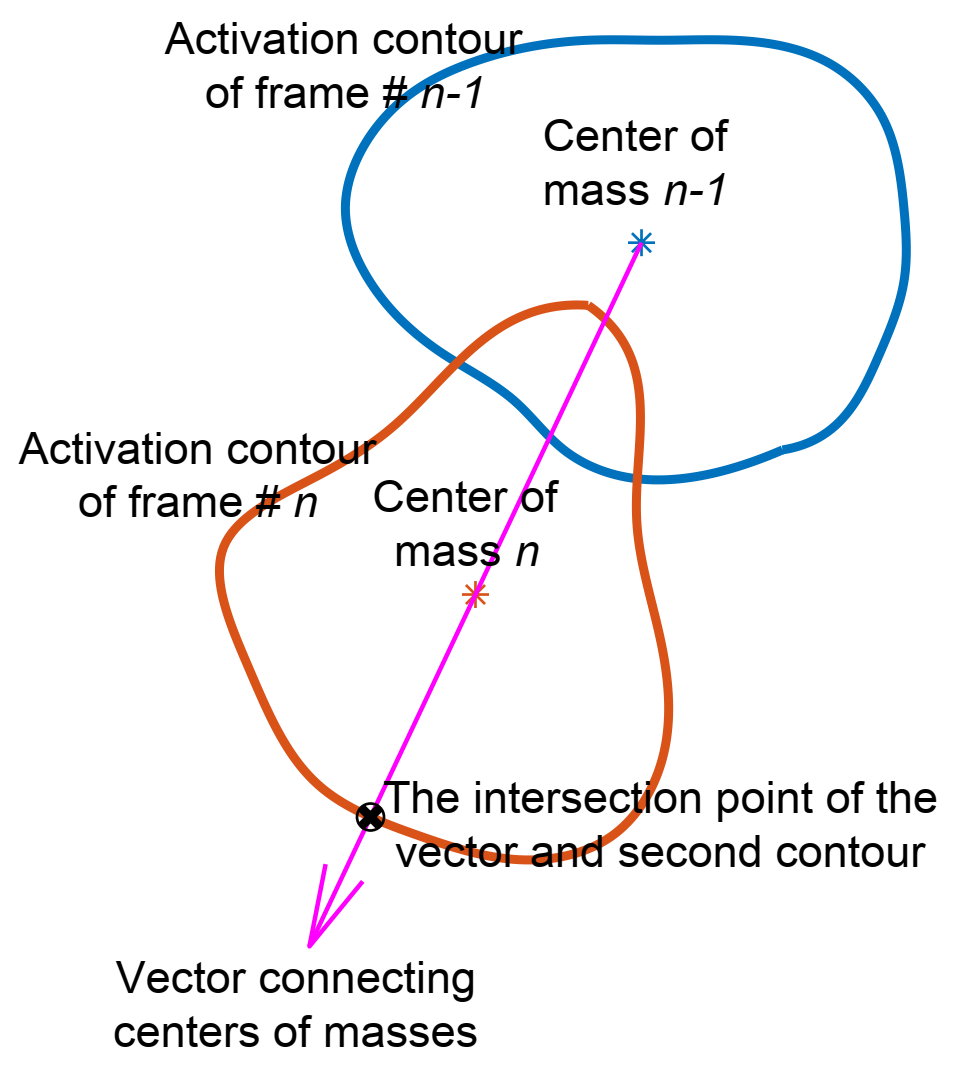

**F**

**Fig. SM2 Estimation of activity onset.** **(A)** Representative curve to explain the method for finding the spontaneous activity onset. In this example, a VSD signal during 340ms (167ms before and after the peak) is shown in blue. Local minima (black asterisks) are first found and a cubic polynomial curve is fitted to it (magenta line). The onset (shown with red asterisk) is then defined as 1 frame prior to the intersection between the original signal and its fitted curve. **(B)** Percent distribution of the difference between the estimated activity onset and true stimulus onset for forelimb evoked activity. A value of difference close to 0 indicates accurate estimation. **(C**-**E)** are similar to **(B)** but for hindlimb, visual, and auditory evoked activities respectively. **(F)** Schematic of the activity trajectory estimation for sensory evoked responses and spontaneous activity motifs. For each two consecutive frames (*n-1* and *n*), activation contours and consequently their centers of masses are found. The direction of spread of activity is determined as the vector connecting both centroids. The next point in the trajectory path (black asterisk) is the intersection of direction of activity (magenta arrowed line) with the activation contour of frame *n* (red contour).

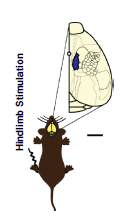

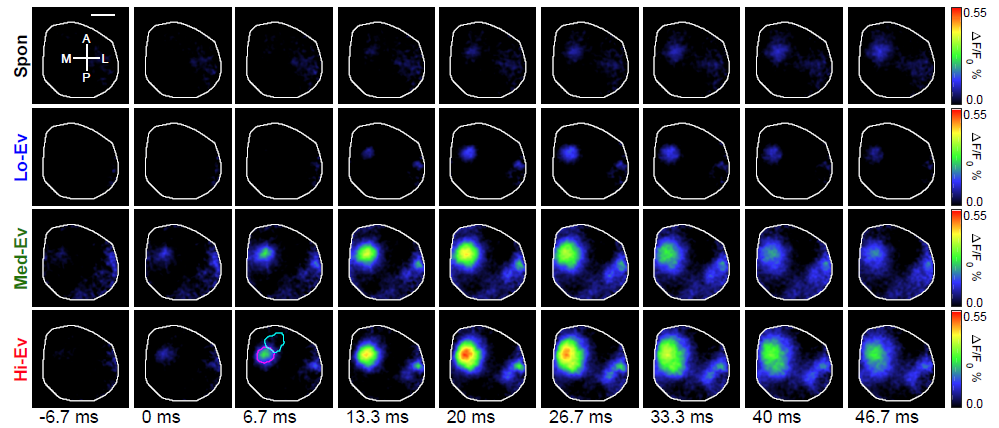

**A**

Anesthetized mouse

Voltage-sensitive dye imaging of cortical activity hemisphere

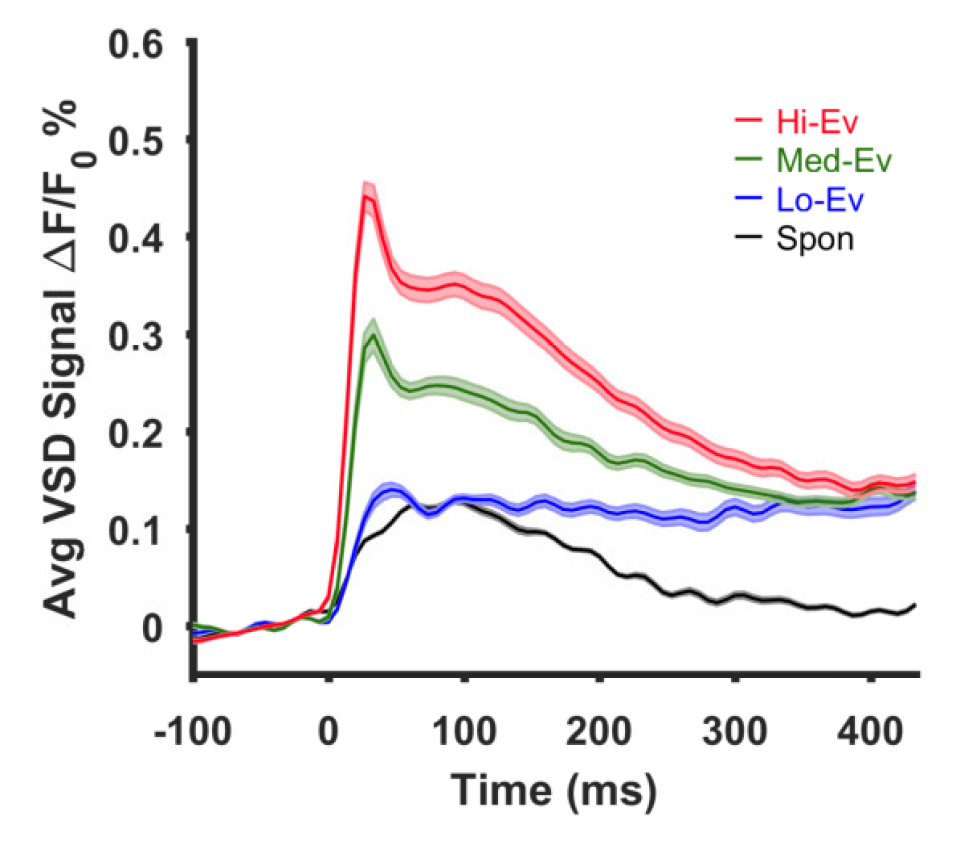

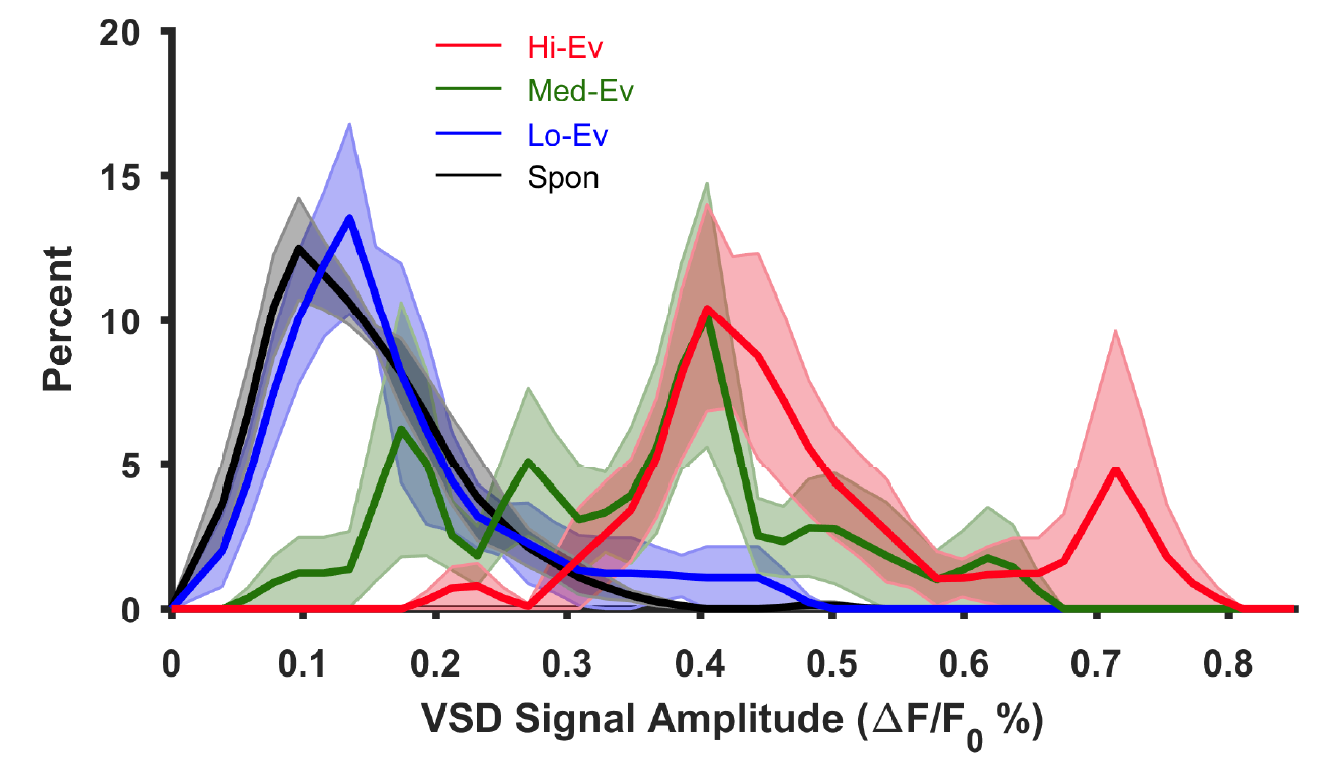

**B**

**C**

**E**

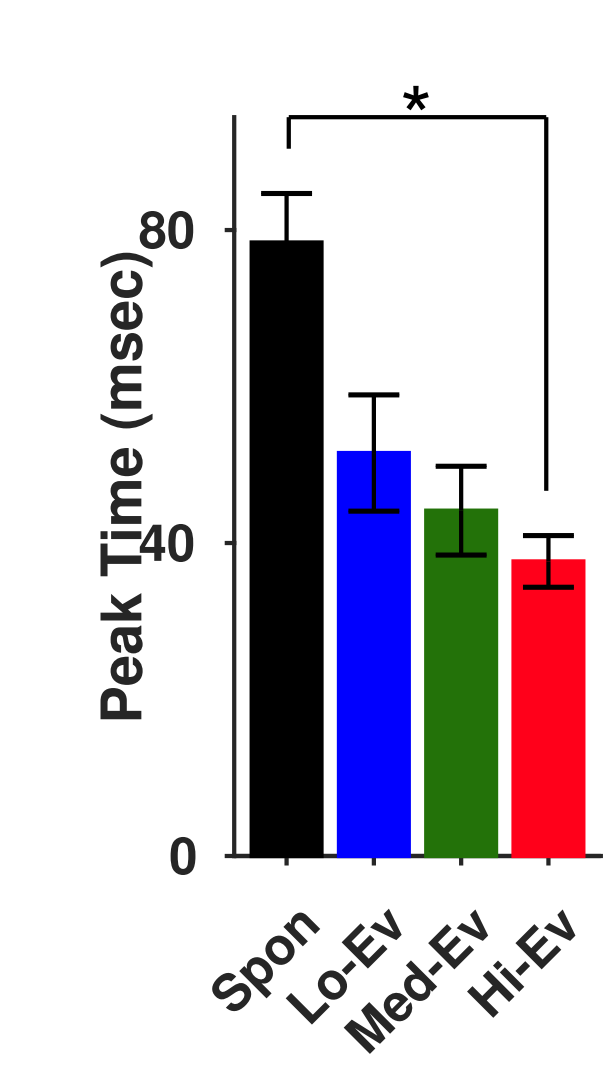

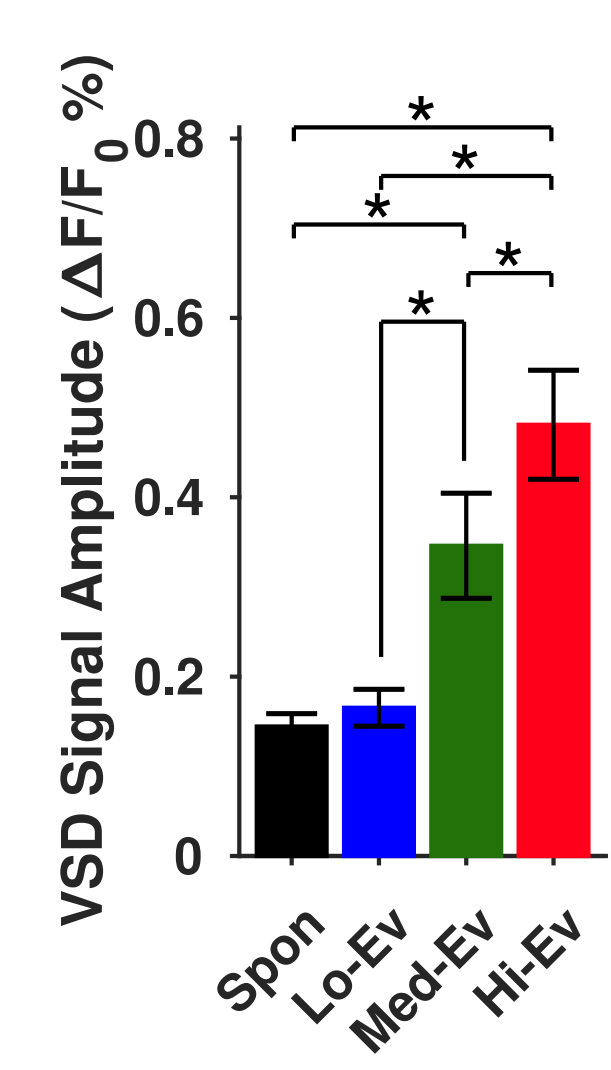

**D**

**Fig. S1 Amplitude of hindlimb-evoked activity is larger than that of spontaneous activity in anesthetized mice**. **(A)** Experimental paradigm (left) and montages of VSD imaging (right). Montage top row shows a representative average of hindlimb spontaneous activity motif. The bottom three rows show the average sensory-evoked cortical activity in response to contralateral hindlimb stimulation with low (Lo), medium (Med), and high (Hi) stimuli strengths. Primary forelimb and hindlimb regions of interest (FLS1 and HLS1 ROIs) are outlined in the third column of the bottom row. Compass lines indicate anterior (A), posterior (P), medial (M) and lateral (L) directions. Scale is 1 mm. **(B)** Plots of the average VSD signal in the HLS1-ROI (n=5 animals) for hindlimb stimulation with different stimulation intensity and spontaneous activity. The curves here represent average over animals calculated from averages over trials for individual animals. Thick lines indicate the mean while shaded regions indicate SEM over animals. 0ms indicates onset of activity. **(C)** Average distributions of VSD signal amplitudes in the HLS1-ROI from 5 animals (normalized to the percentage of occurrences). VSD Signal Amplitude = (peak ΔF/F_0_) – (mean of baseline) as depicted in (Fig. 1B). Shaded regions show SEM over animals. **(D-E)** Mean±SEM values of VSD signal amplitudes and time-to-peak respectively. * indicates p<0.05, repeated measure ANOVA with post-hoc Tukey-Kramer correction.

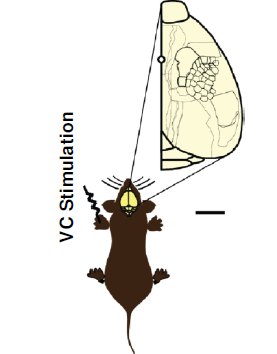

**A**

**B**

Anesthetized

Voltage-sensitive dye imaging of right cerebral hemisphere

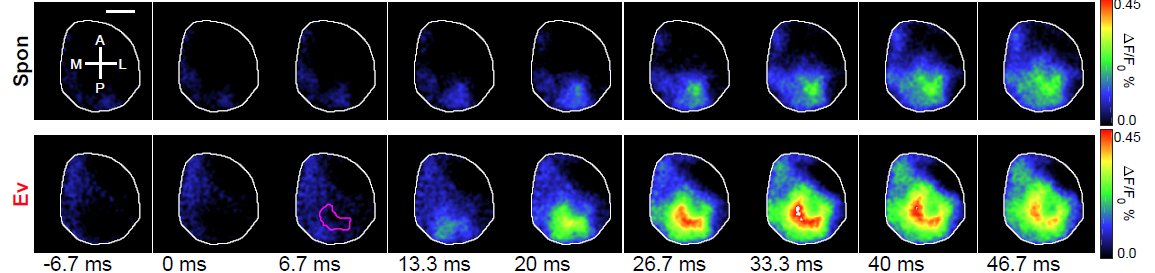

**C**

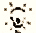

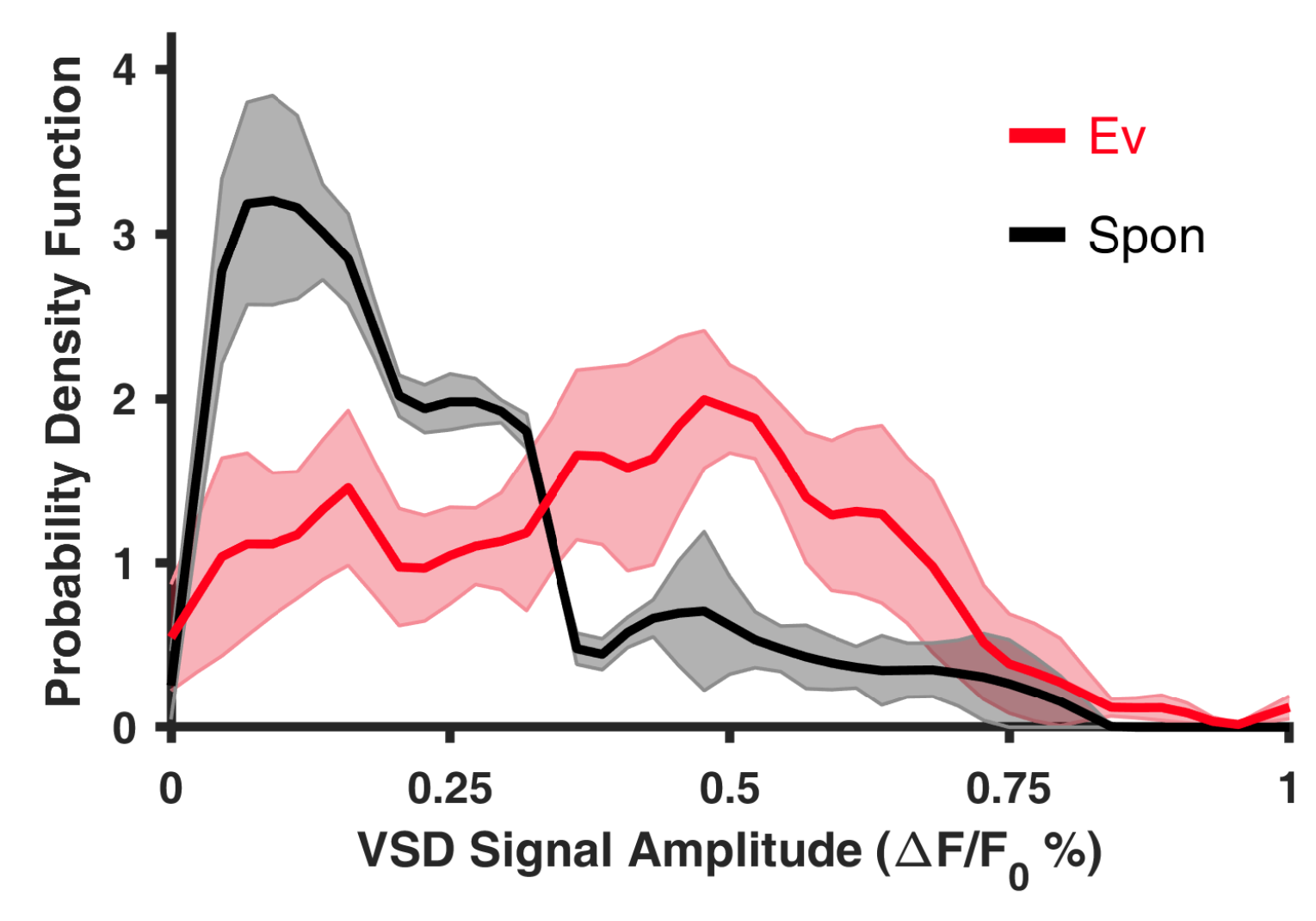

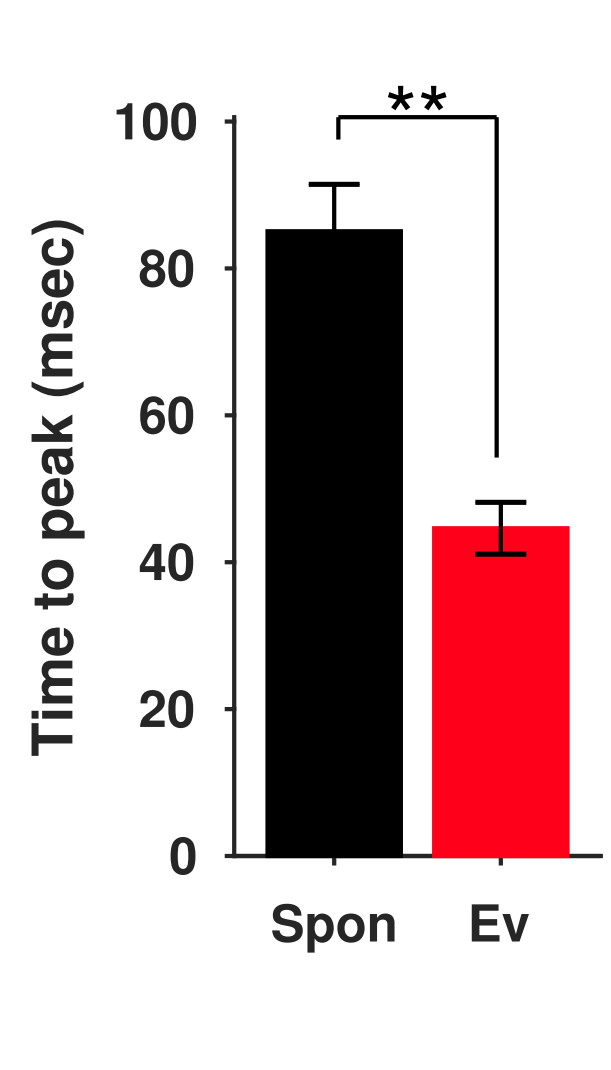

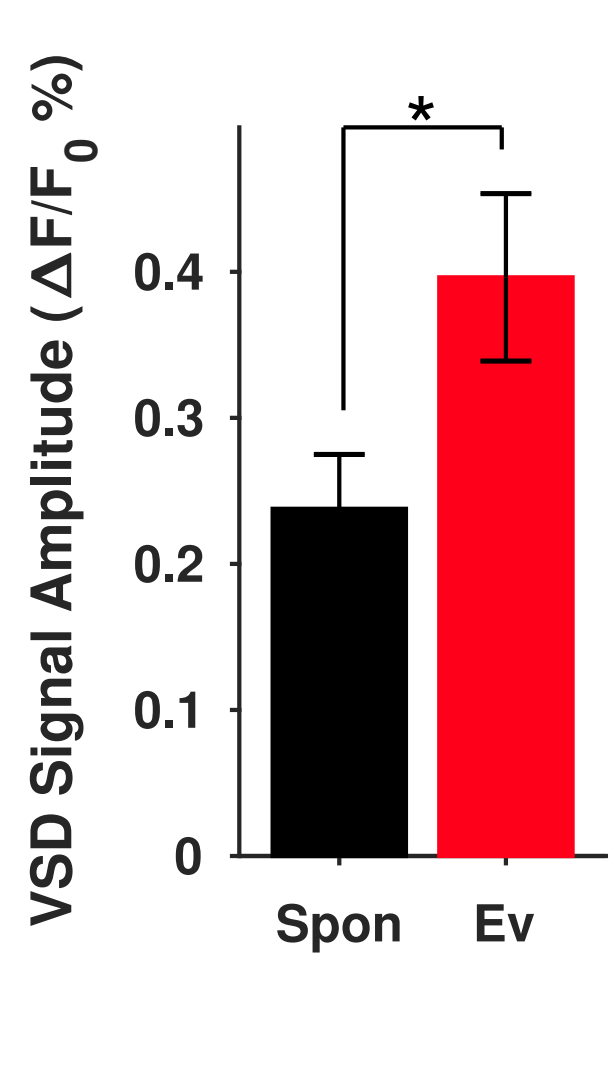

**E**

**D**

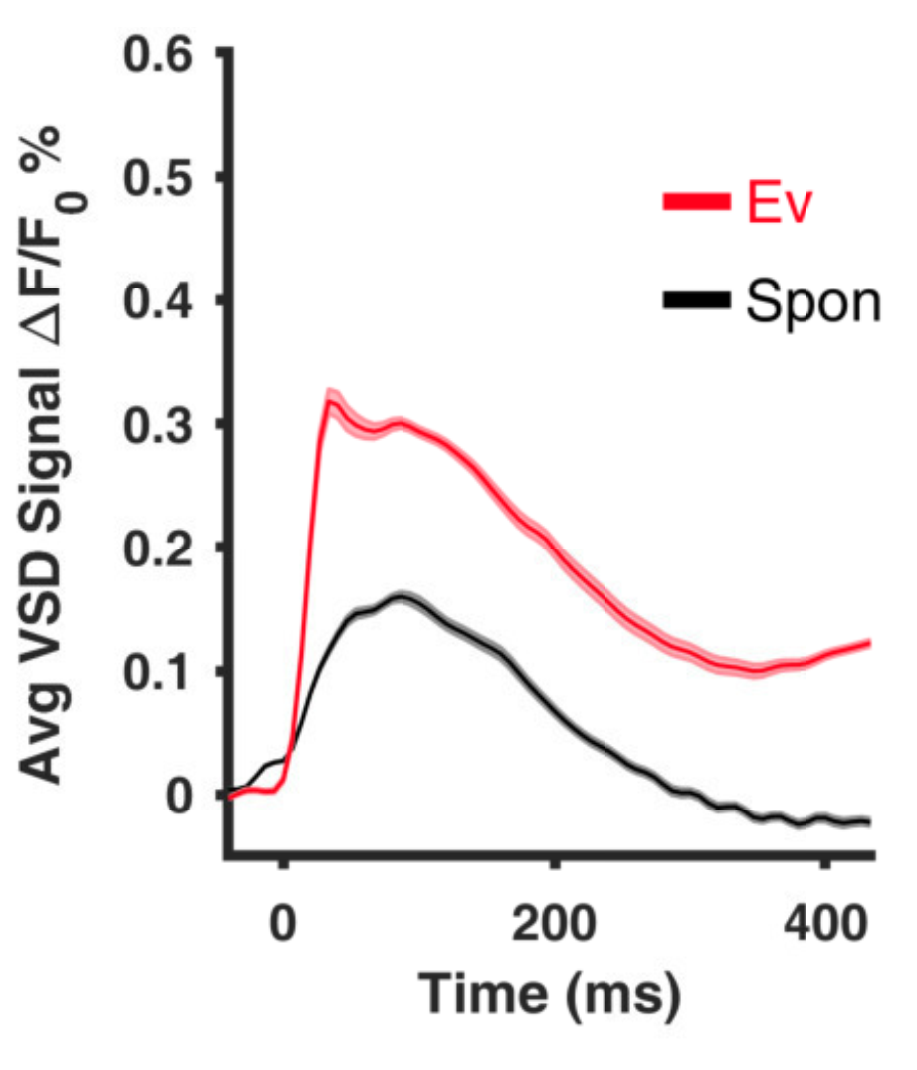

**Fig. S2 Amplitude of visual-evoked activity is larger than that of spontaneous activity motifs in anesthetized mice**. **(A)** Experimental paradigm (left) and montages of VSD imaging (right). Montage top row shows a representative average visual motifs in spontaneous activity. The bottom row shows the average evoked activity in response to contralateral visual stimulation. Primary visual region of interest (VC ROI) is outlined in the third column of the bottom row. Compass lines indicate anterior (A), posterior (P), medial (M) and lateral (L) directions. Scale is 1 mm. **(B)** Plots of the average VSD signal in the VC-ROI (n=4 animals). Thick lines indicate the mean while shaded regions indicate SEM over animals **(C)** Average distributions of VSD signal amplitudes in the VC-ROI from 4 animals (normalized to the percentage of occurrences). VSD Signal Amplitude = (peak ΔF/F_0_) – (mean of baseline) as depicted in (Fig. 1B). Shaded regions show SEM over animals. **(D-E)** Mean±SEM values of VSD signal amplitudes and time-to-peak respectively. * and ** indicate p<0.05 and p<0.01 respectively, paired sample t-test.

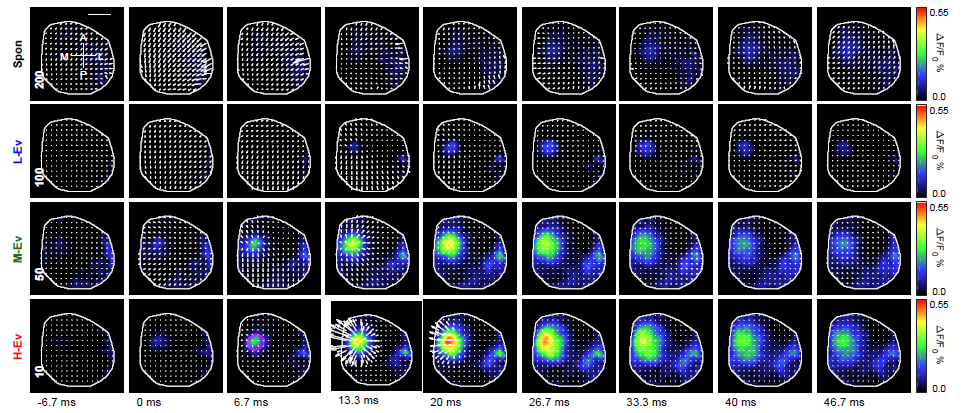

**A**

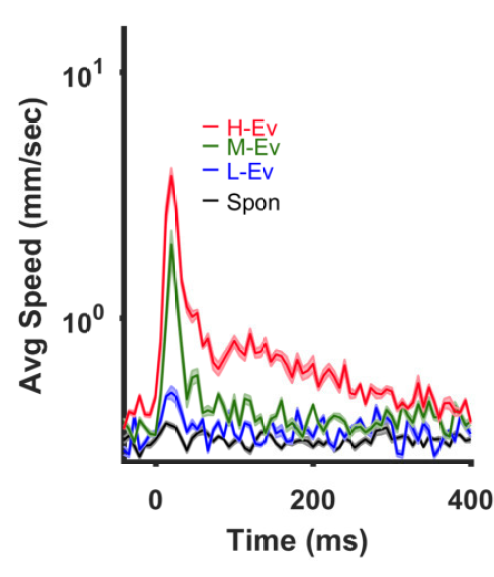

**B**

**C**

**D**

**E**

**F**

HLS1 ROI

HLS1 ROI

FLS1 ROI

PTA ROI

HLS1 ROI

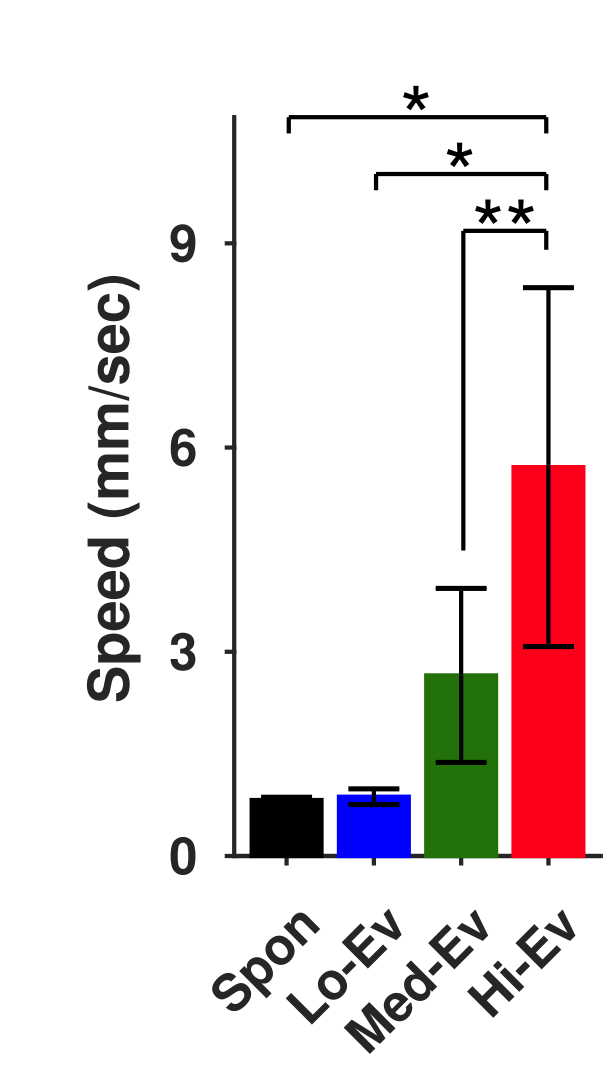

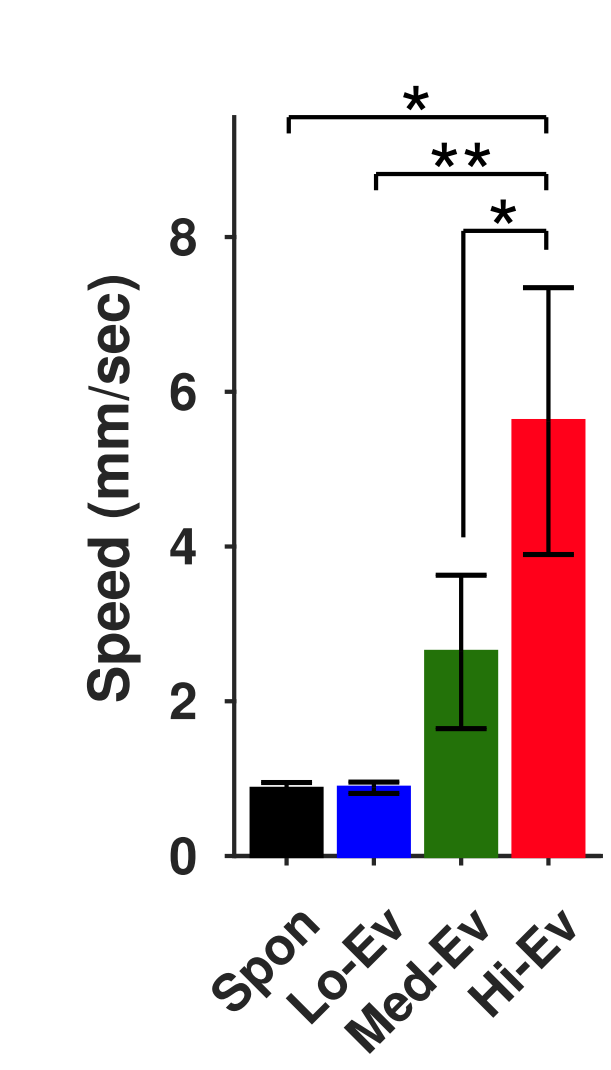

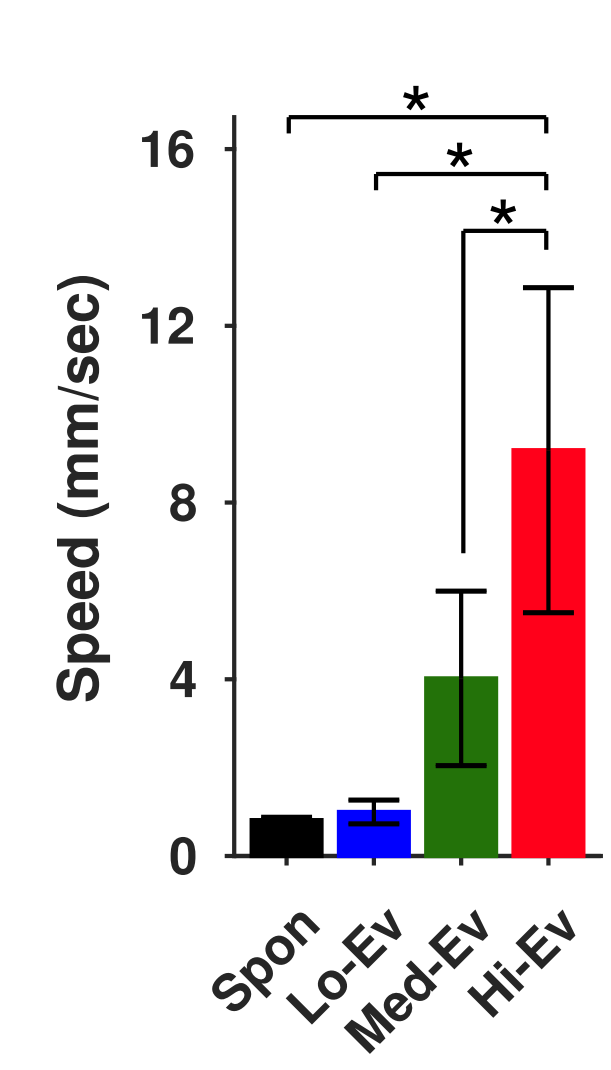

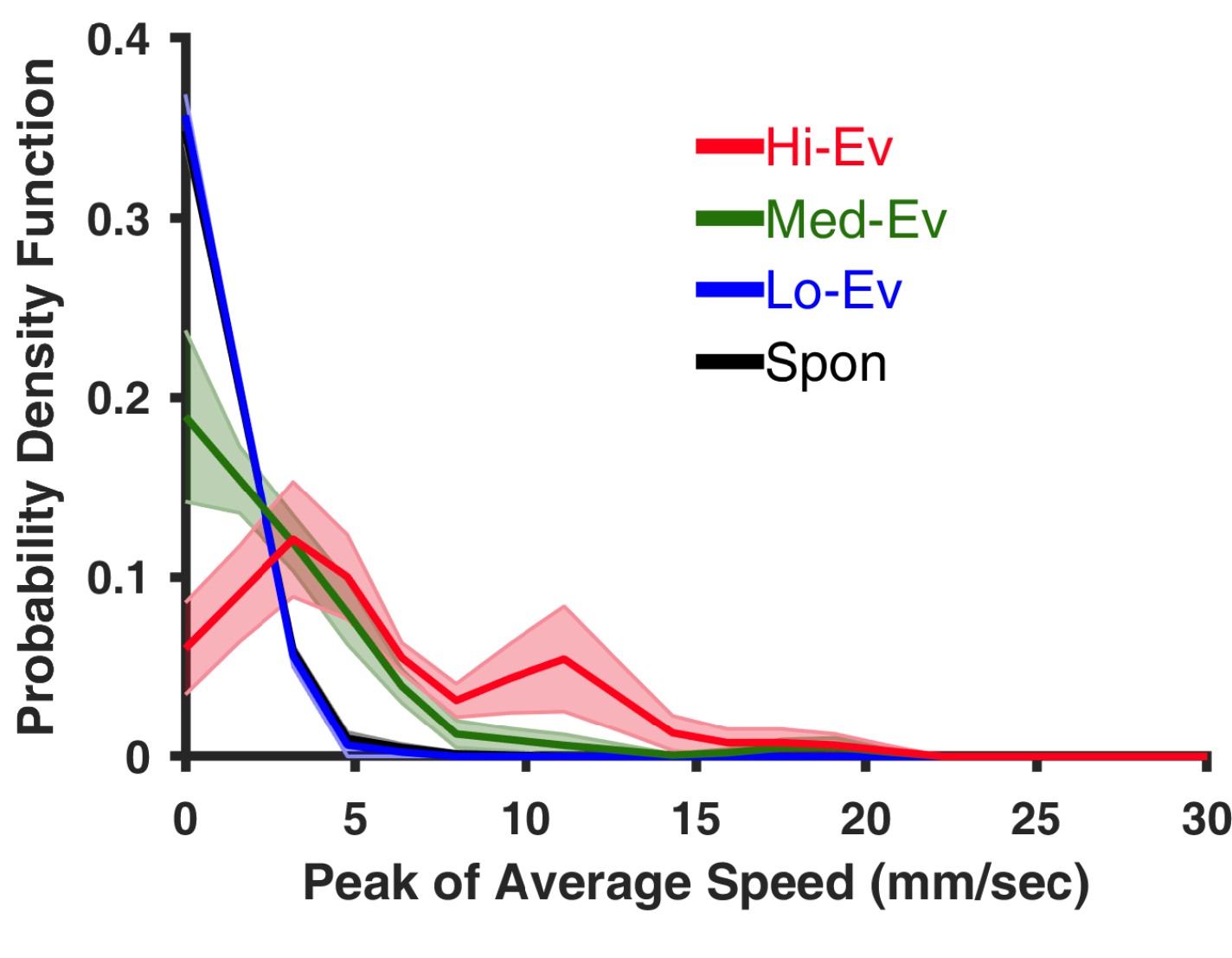

**Fig. S3 Propagation speed of hindlimb-evoked activity is larger than that of spontaneous activity in anesthetized mice**. **(A)** Montage top row shows a representative average motif of hindlimb spontaneous activity motifs overlaid with the velocity vector field determined by optical flow analysis. The bottom three rows show similar montages for average evoked (Ev) activity in response to contralateral hindlimb stimulation with low (Lo), medium (Med), and high (Hi) stimuli strengths. HLS1 ROI is outlined in the third column of the bottom row. Compass lines indicate anterior (A), posterior (P), medial (M) and lateral (L) directions. Scale is 1 mm. Numbers in the first column indicate scale factor for drawing velocity vector fields. **(B)** Plots of the average speed signal in the HLS1-ROI (n=5 animals). The summary graphs represent average over animals. Note that the scale in the y axis is logarithmical. Thick lines indicate the mean while shaded regions indicate SEM over animals. **(C)** Average distributions of peak average speed in the HLS1-ROI (normalized to the percentage of occurrences). Shaded regions show SEM over animals. **(D**-**F)** Mean±SEM values of peak of average speeds in the HLS1, FLS1, and PTA ROIs respectively. Note that different y scale is used for each ROIs. * and ** indicate p<0.05 and p<0.01 respectively, repeated measure ANOVA with post-hoc Tukey-Kramer correction.

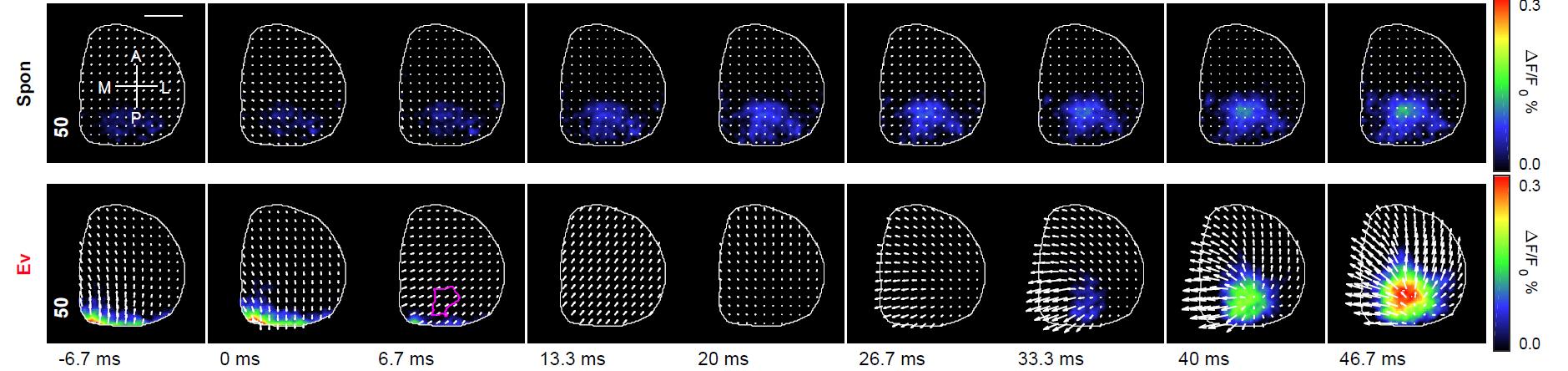

**A**

**B**

**C**

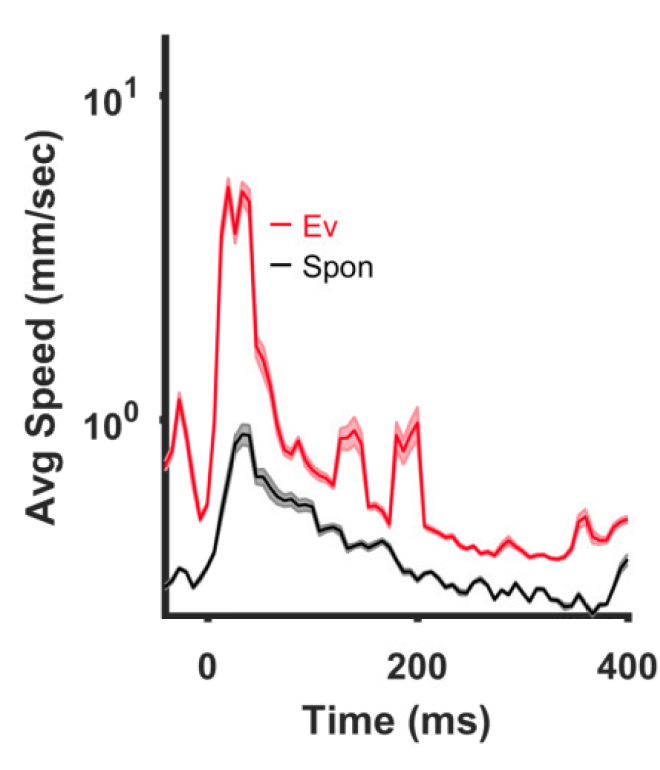

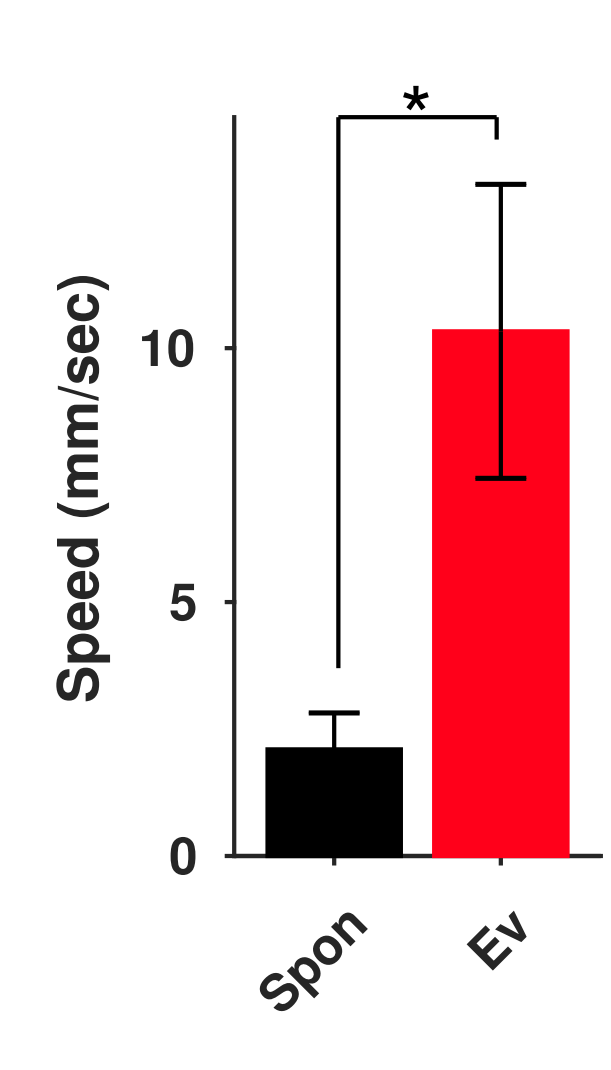

**D**

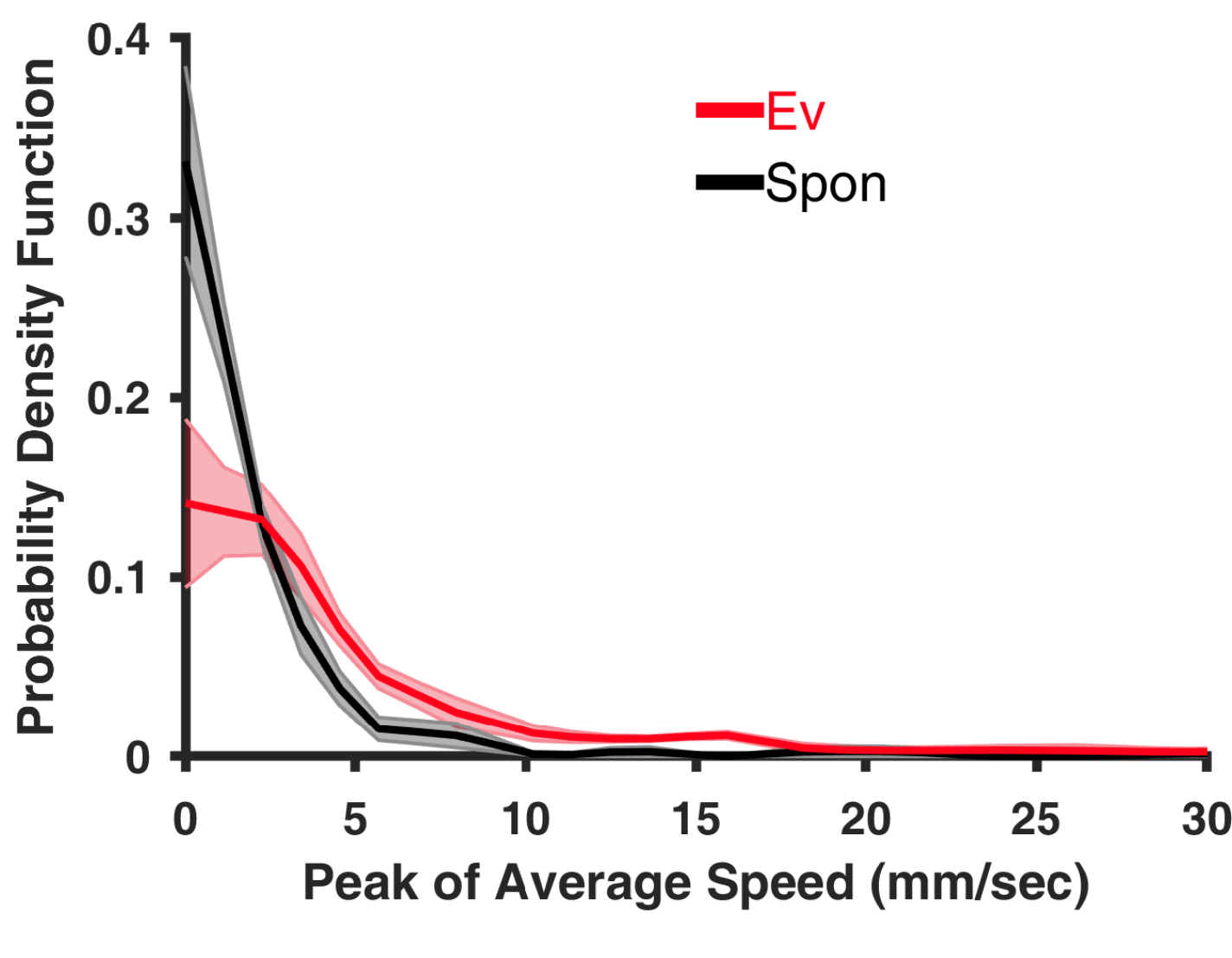

**Fig. S4 Propagation speed of visual-evoked activity is larger than that of spontaneous activity in anesthetized mice**. **(A)** Montage top row shows a representative average motif of visual spontaneous activity motifs overlaid with the velocity vector field determined by optical flow analysis. The bottom row shows a similar montage for average evoked activity in response to contralateral visual stimulation. VC ROI is outlined in the third column of the bottom row. Compass lines indicate anterior (A), posterior (P), medial (M) and lateral (L) directions. Scale is 1 mm. The numbers in the first column indicate the scale factor for drawing the velocity vector fields. **(B)** Plot of the average speed signal in the VC-ROI (n = 4 animals). The summary graphs represent average over animals. Note that the scale in the y axis is logarithmical. Thick lines indicate the mean while shaded regions indicate the SEM over animals. **(C)** Average distributions of peak average speed in the VC-ROI (normalized to the percentage of occurrences). Shaded regions denote the SEM over animals. **(D)** Mean±SEM values of peak of average speed in the VC ROI. * indicate p<0.05, paired sample t-test.

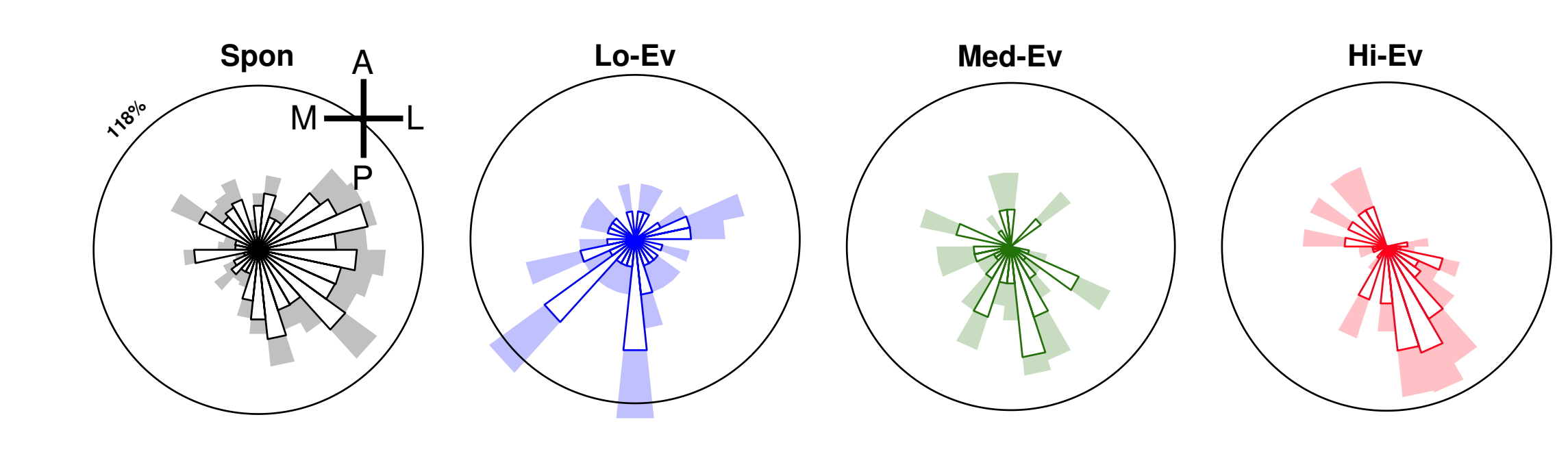

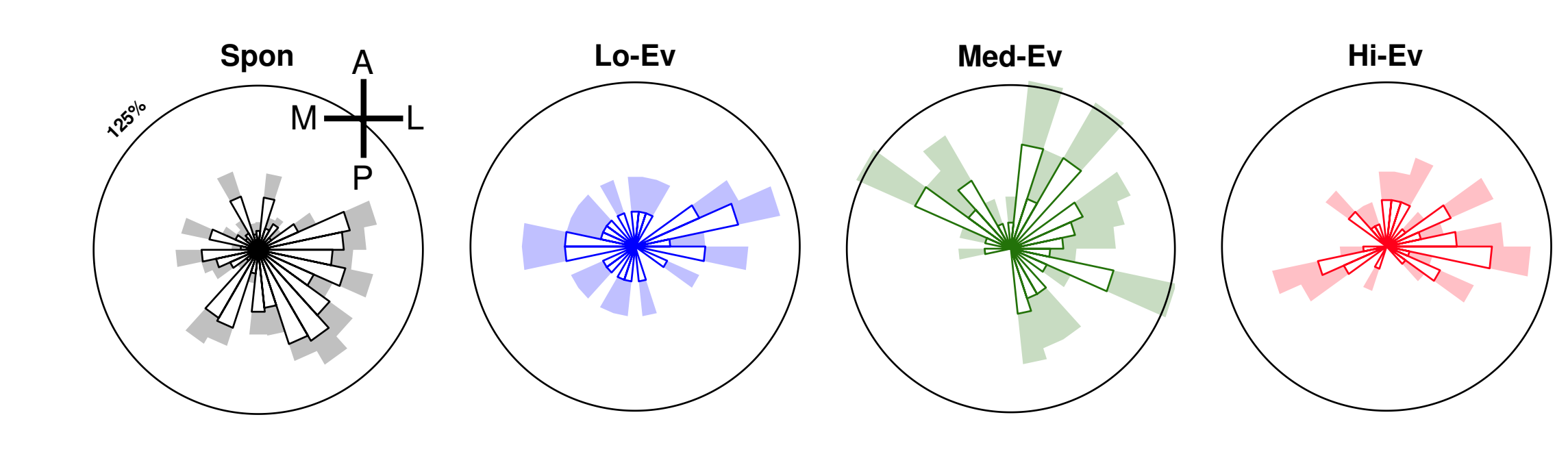

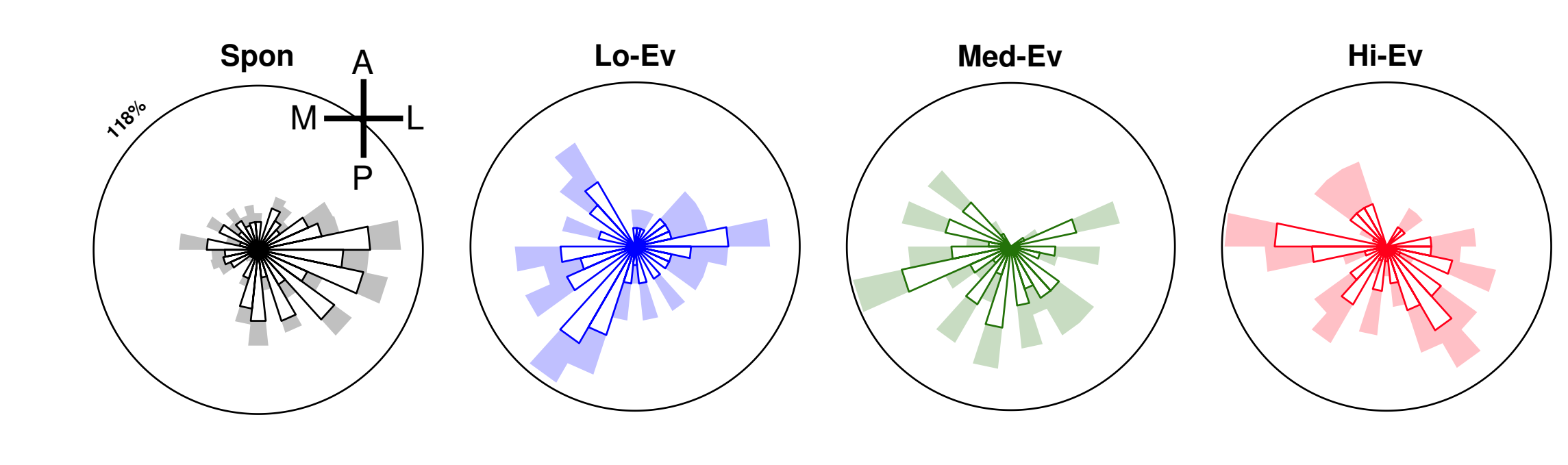

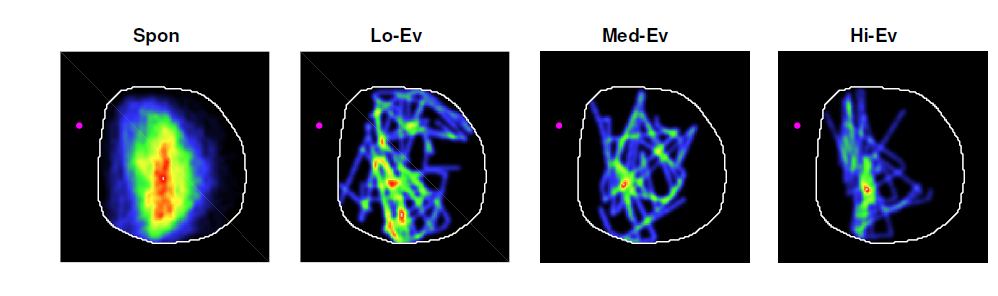

**E**

**H**

Animal # 5

**HLS1 ROI**

**A**

**FLS1 ROI**

**C**

**B**

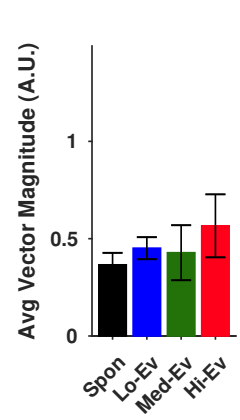

**D**

**PTA ROI**

**G**

**F**

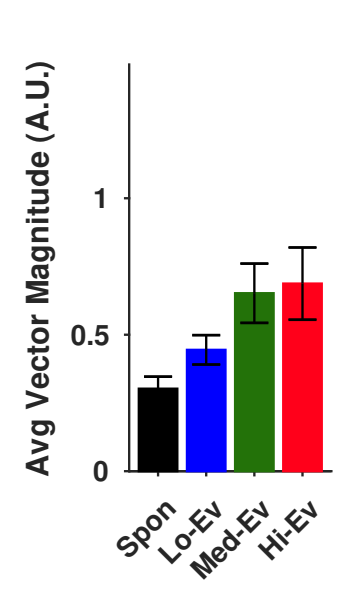

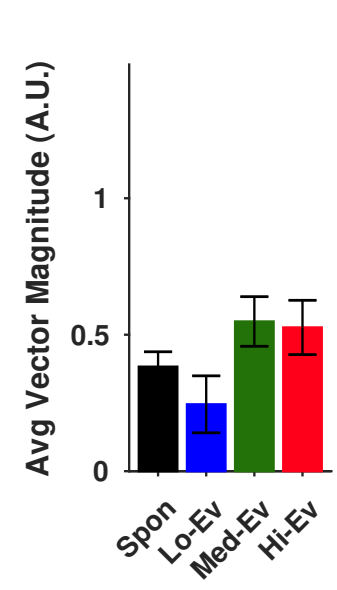

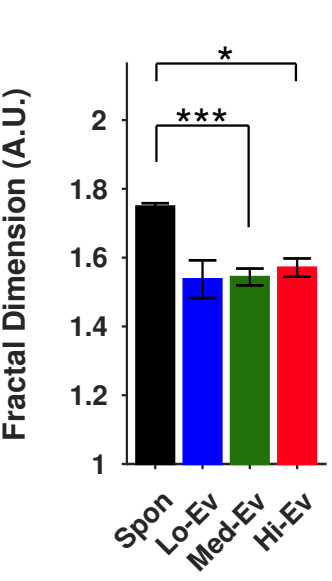

**Fig. S5 Propagation patterns of hindlimb-evoked activity converge as stimulus levels increase and are less complex than those of spontaneous activity in anesthetized mice**. **(A)** Average distributions of the directions of the peak velocity vectors for spontaneous (Spon) activity and evoked (Ev) activity elicited with Lo, Med, and Hi stimulus levels in the HLS1 ROI over all animals (n=5). The values are normalized to the percentage of occurrences. **(B)** Magnitude of the average of normalized velocity vectors. Error bars represent the SEM over animals. **(C**-**D)** and **(E**-**F)** similar to (**A**-**B**) but for FLS1 and PTA ROIs respectively. **(G)** Normalized histogram of activity trajectories represented as a heat map with cold and warm colors indicating smaller and larger numbers respectively of activity passing through a given point on the cortical surface. **(H)** Mean±SEM values of the Hausdorff fractal dimension of the heat maps. * and *** indicate p<0.05 and p<0.001 respectively, repeated measure ANOVA with post-hoc Tukey-Kramer correction.

**A**

**VC ROI**

**B**

Animal # 1

**C**

**D**

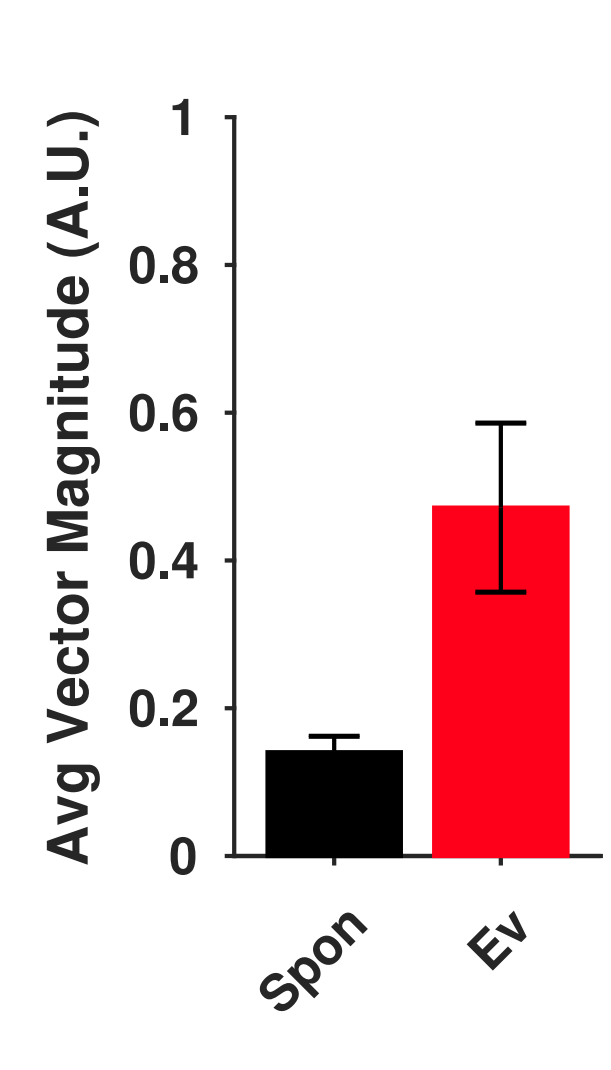

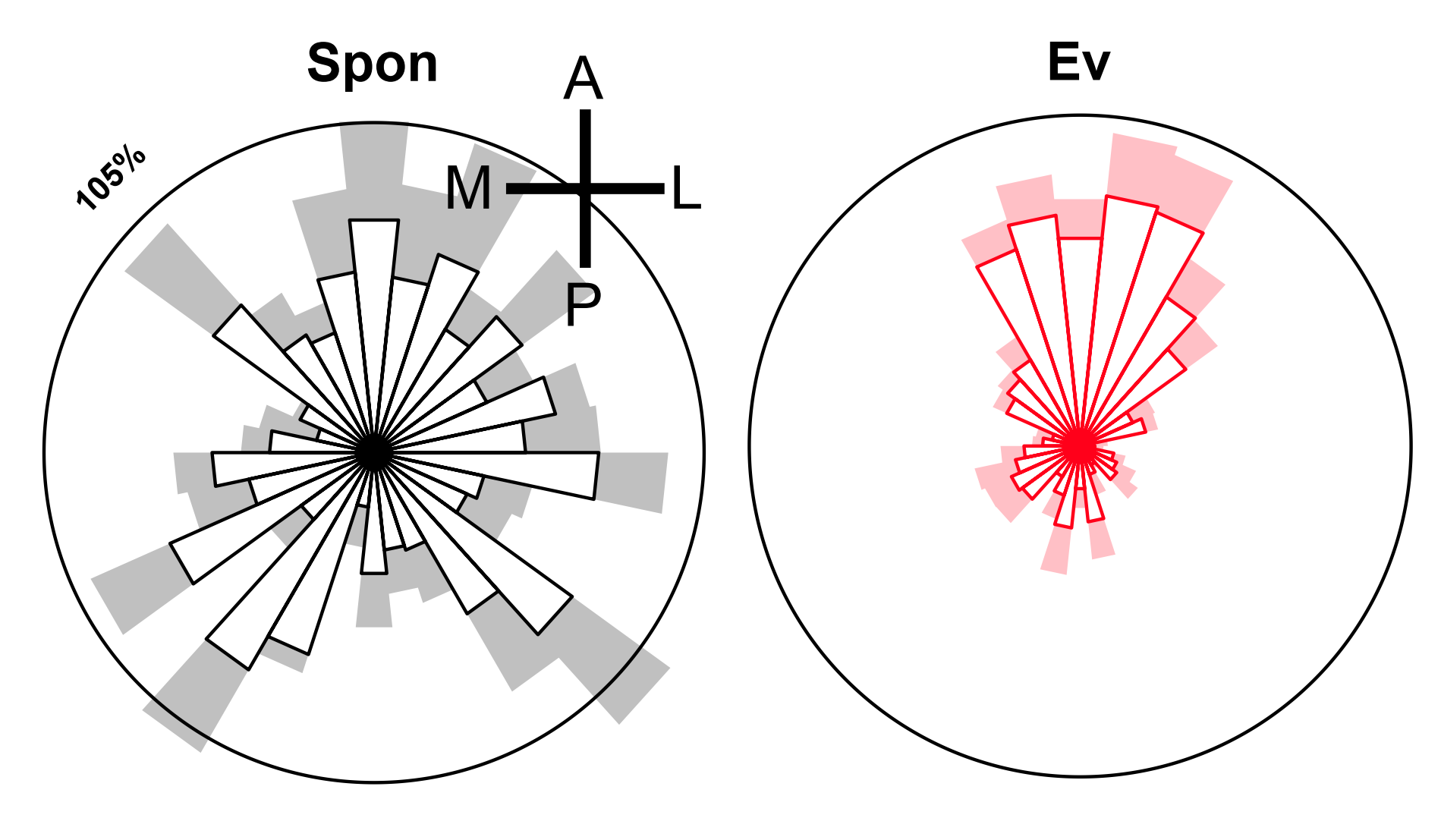

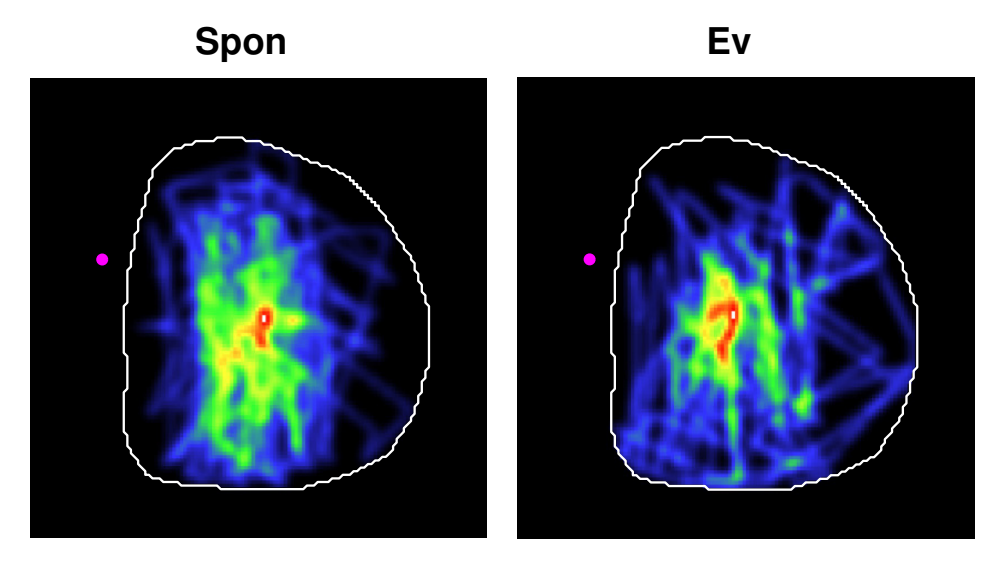

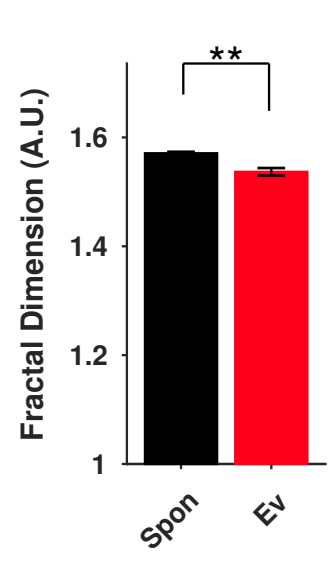

**Fig. S6 Propagation patterns of visual-evoked activity are less complex than those of spontaneous activity in anesthetized mice**. **(A)** Average distributions over animals (n=4) of the directions of the peak velocity vectors in the VC ROI. Distributions averaged over animals and normalized to the percentage of occurrences are represented for spontaneous (Spon) activity and evoked (Ev) activity elicited with only one stimulus level. **(B)** Magnitude of the average of normalized velocity vectors. **(C)** Normalized histogram of activity trajectories represented as a heat map with cold and warm colors indicating smaller and larger numbers of activity passing through a given point on the cortical surface respectively. **(D)** Mean±SEM values of the Hausdorff fractal dimension of the heat maps. ** indicate p<0.01, paired t-test.

**HL Spon**

**VC Spon**

**B**

**C**

**A**

**D**

**E**

**F**

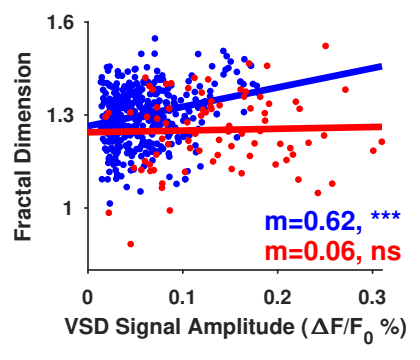

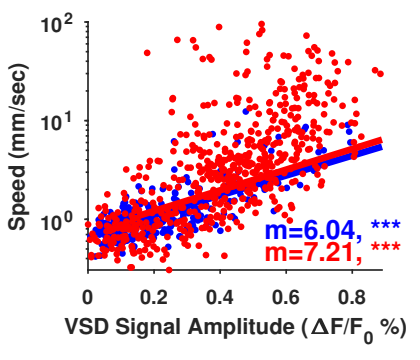

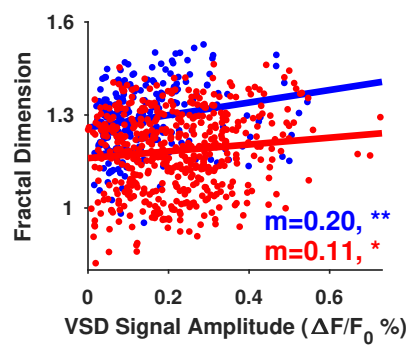

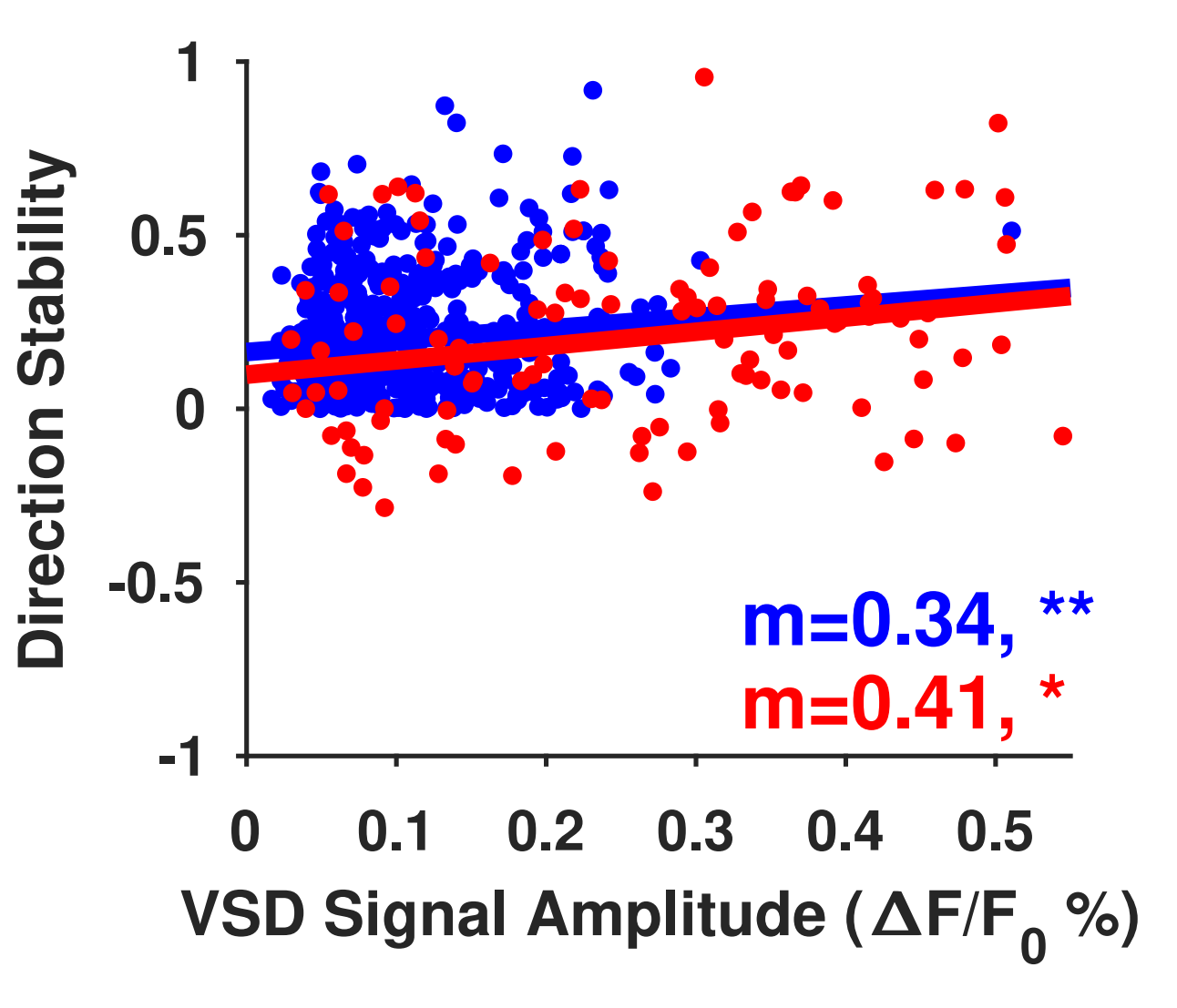

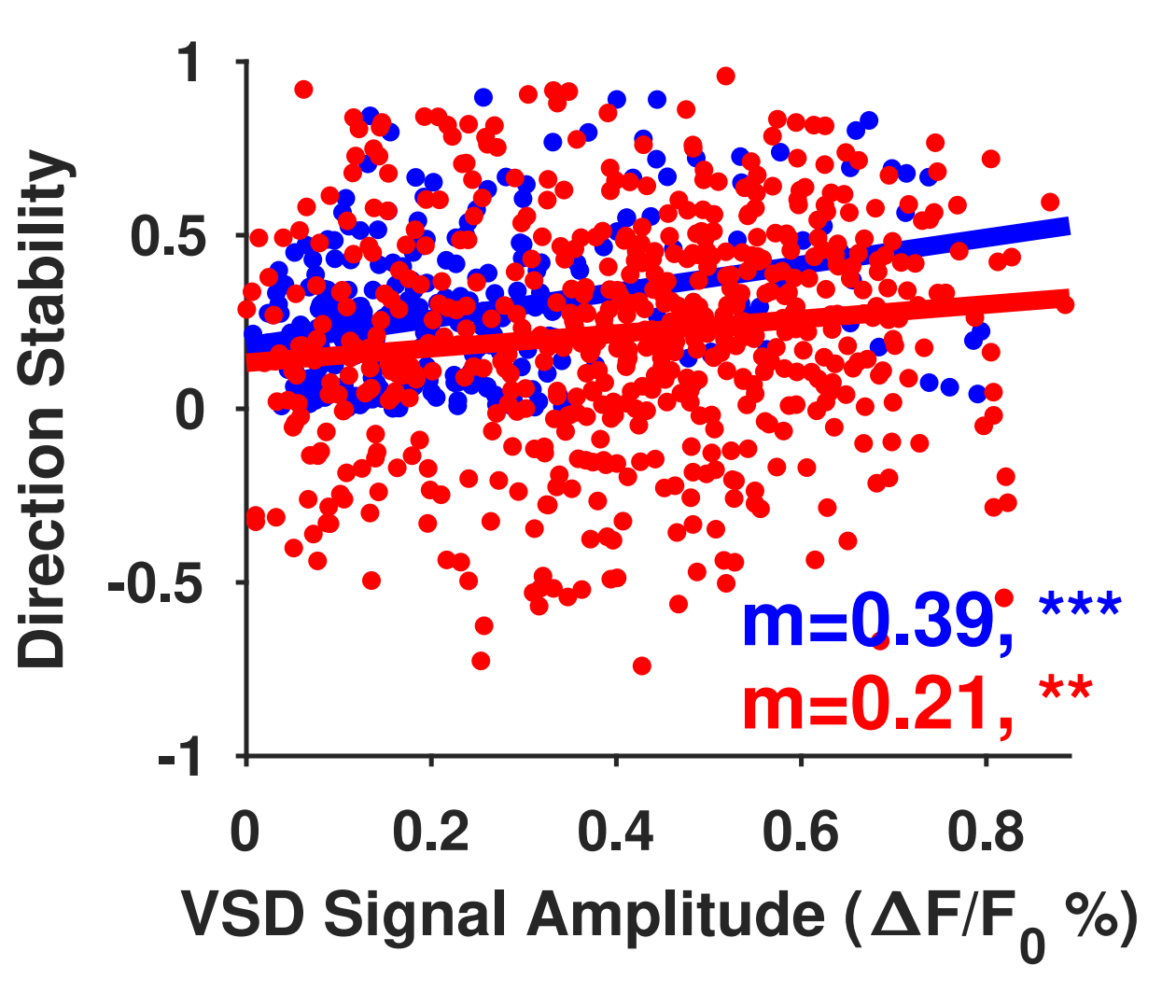

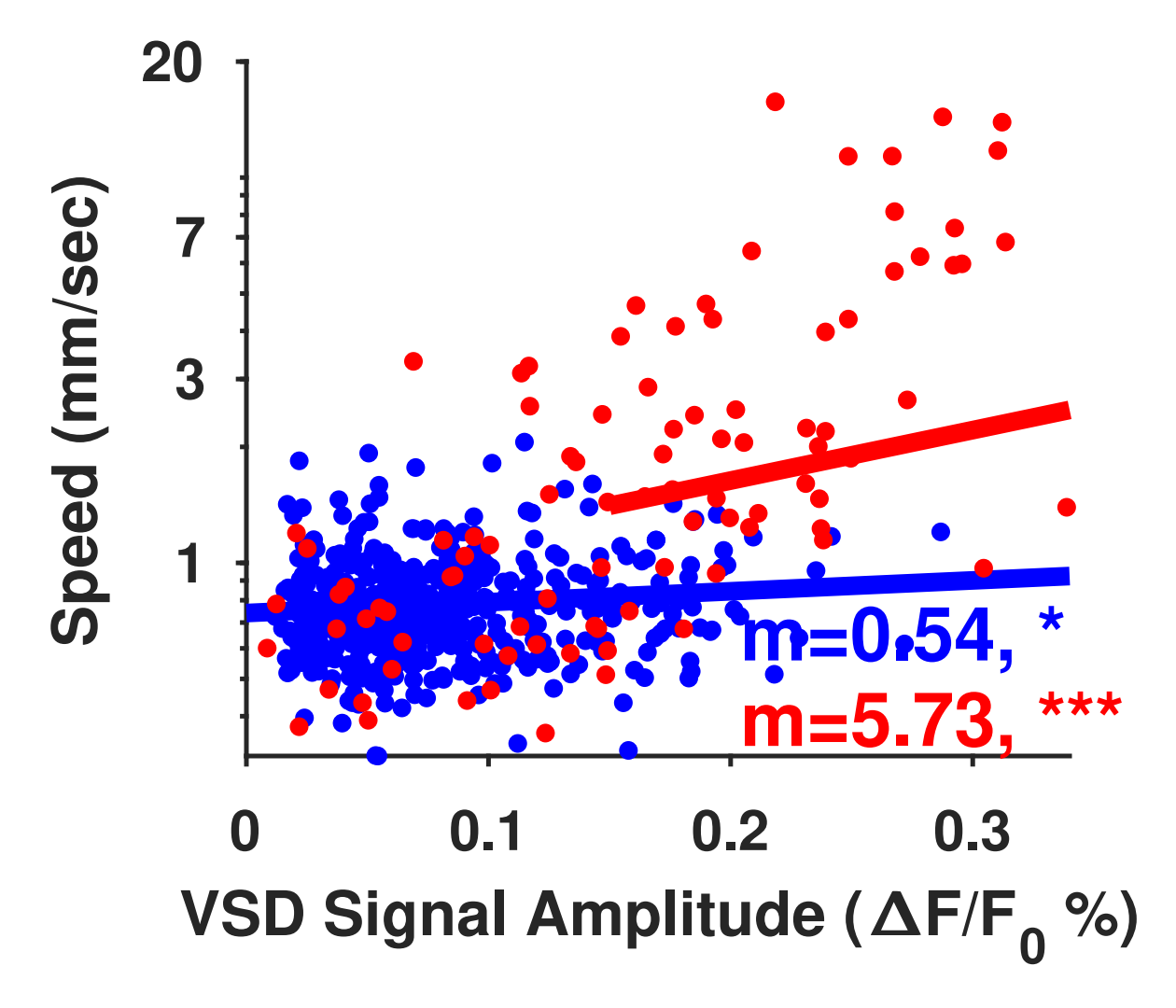

**Fig. S7 Speed, flow direction stability, and trajectory space of spontaneous activity patterns are positively correlated with spontaneous activity amplitude in anaesthetized mice**. **(A)** Scatter plot of VSD signal amplitudes for hindlimb spontaneous motifs (blue) and evoked (red) activity versus peak velocity vectors in the FLS1 ROI for anaesthetized mice. Each dot represents one trial. The data is pooled from all animals (n=5). The lines represent linear regression models fitted to the corresponding data. **(B)** Scatter plot and linear regression fit for VSD signal amplitude for hindlimb spontaneous (blue) motifs and evoked activity (red) in the PTA ROI vs flow direction stability for anesthetized mice. **(C)** Similar to **(A)** but for VSD signal amplitudes in the entire imaging window vs fractal dimensions of spontaneous motifs (blue) and evoked activity (red). (**D-F**) Similar to (**A-C**) but for visual anaesthetized spontaneous motifs in VC, VC, and the entire imaging window ROIs respectively. The data is pooled from all animals (n=4). m is the estimated slope. *, **, and *** indicates p<0.05, p<0.01, and p<0.001 respectively, for the p-value of the linear regression slope. Notice that speed axis in **A** and **D** is logarithmic scale.
